# From heterogeneity to homogeneity: coordination of siderophore gene expression among clonal cells of the bacterium *Pseudomonas aeruginosa*

**DOI:** 10.1101/2021.01.29.428812

**Authors:** Subham Mridha, Rolf Kümmerli

## Abstract

There has been great progress in understanding how bacterial groups coordinate social actions, such as biofilm formation, swarming and public-goods secretion. Less clear, however, is whether the seemingly coordinated responses observed at the group level actually mirror what individual cells do. Here, we use a microscopy approach to simultaneously quantify the investment of individual cells of the bacterium *Pseudomonas aeruginosa* into two public goods, the siderophores pyochelin and pyoverdine. Using gene expression as a proxy for investment, we initially observed no coordination but high heterogeneity and bimodality in siderophore gene expression across cells. With increasing cell density, gene expression became homogenized across cells, accompanied by a shift from pyochelin to pyoverdine expression. We found positive correlations in the expression of pyochelin and pyoverdine genes across cells, and show that cell-to-cell variation is driven by differences in cellular metabolic states. We propose a model explaining how variation in internal iron stocks can spur initial erratic gene expression, while siderophore-mediated signalling and intra-cellular feedbacks later on can induce highly coordinated gene expression and synchronized shifts from pyochelin to pyoverdine. Our work provides new insights into bacterial collective decision-making processes and reveals a three-phase chronobiological siderophore investment cycle in *P. aeruginosa*.

## INTRODUCTION

Bacteria perform many actions that are vital for survival and growth outside their cells [1]. For example, bacteria secrete enzymes to digest extra-cellular polymers and proteins, siderophores to scavenge iron, and biosurfactants to enable swarming on wet surfaces [2–5]. A consequence of molecule secretion is that the benefit of the performed actions can accrue to other cells, because digested nutrients, iron-loaded siderophores and biosurfactants become accessible to other individuals within a group [6–9]. Because of their group-level effects, molecule production often occurs in a coordinated fashion, relying on molecular mechanisms such as quorum sensing and other signaling circuits, integrating information across individuals within a group [10–12]. There has been tremendous interest in understanding the molecular basis of these regulatory circuits, and the ecological and evolutionary consequences of coordinated molecule production [5, 13–15]. Yet, many of the insights gained on group-coordinated behavior originate from batch-culture experiments, where behavioral responses were averaged across millions of cells. In contrast, we still know little on how individual cells behave within a group, and whether decisions taken by individuals indeed match the patterns we interpret as coordinated group response at the population level [16–18].

Here, we tackle these open issues by studying single-cell social behavior in the opportunistic human pathogen *Pseudomonas aeruginosa* PAO1. This species produces a suit of extra-cellular compounds that can benefit other group members [10]. In our study, we focus on the investment strategies of individual cells into the two iron-chelating siderophores pyoverdine and pyochelin. Siderophores are produced and secreted in response to iron limitation in order to scavenge this essential nutrient from the environment [19, 20]. Siderophore-iron complexes are subsequently taken up by cells via cognate receptors [21]. Due to their extra-cellular mode of action and their high diffusivity, siderophores can be considered ‘public goods’, benefiting other cells in a clonal group [5].

*P. aeruginosa* produces two siderophores, pyochelin and pyoverdine via non-ribosomal peptide synthesis [22]. The regulation of these two siderophores involves three levels. The first level is mediated by Fur (ferric uptake regulator). Fur blocks siderophore synthesis, but loses its inhibitory effect when intra-cellular iron stocks become depleted [20, 23–25]. Fur de-repression initiates basal siderophore gene expression, with evidence existing that de-repression happens earlier for pyochelin than for pyoverdine [26]. The second level involves a membrane-embedded signaling cascade, where incoming siderophore-iron complexes trigger a positive feedback loop that increases siderophore production [27, 28]. The third level is pleiotropic in nature. It is based on a hierarchical regulatory linkage between the two siderophores, whereby pyoverdine suppresses pyochelin production within cells based on a yet unknown mechanism [26, 29].

All the three levels interact and there is likely heterogeneity in the relative strength of each level across cells in a clonal population. For one thing, cells might vary in their intracellular iron stocks, which affects the strength of Fur repression/de-repression [23] and in turn could determine whether a cell primarily invests in pyochelin, pyoverdine, or none of the siderophores [26]. Furthermore, the extracellular iron availability and local cell density each individual experiences likely varies between cells, which will influence the uptake rate of siderophore-iron complexes and thus the strength of the positive feedback loops. Moreover, it is reasonable to assume that the regulatory mechanisms underlying the three input levels are intrinsically noisy themselves [17]. Given all these sources of variation, a key question is how groups of cells can coordinate and fine-tune their siderophore production strategy. One extreme answer could be that there is no coordination and that the observed population level responses [30–32] might merely be the sum of its heterogenous individual members. Conversely, it could be that the regulatory circuits operate in a way that fosters specialization [33], where fractions of cells in a population invest either in pyochelin or pyoverdine.

Here, we examine these possibilities by using single-cell microscopy to simultaneously track the investment of individual cells into pyochelin and pyoverdine, across a range of media differing in iron availability, and across time following a growth cycle from low to high cell density. We used the expression of pyochelin and pyoverdine synthesis genes as proxies for siderophore investment levels. Specifically, we grew bacteria in batch cultures for 24 hours in the different media, and extracted small samples from these cultures at 3-hour intervals to quantify the expression of pyochelin and pyoverdine genes in cells using fluorescent gene-reporter fusions. Crucially, we constructed a double-fluorescent gene reporter, which allowed us to simultaneously obtain investment proxies for both siderophores for individual cells, and to establish gene expression correlation patterns across cells within populations.

## MATERIALS AND METHODS

### Strains and strain construction

We used the standard laboratory strain *P. aeruginosa* PAO1 (ATCC 15692) for all our experiments, which produces the siderophores pyoverdine and pyochelin [26, 34]. In the genetic background of this wild type strain, we chromosomally integrated fluorescent gene reporter constructs to quantify siderophore gene expression. All constructs were integrated at the *attTn7* site using the mini-Tn7 system [35]. We used the following single-gene reporter fusions previously constructed by Rezzoagli et al. [36]: PAO1*pvdA*:*mcherry*, in which the promoter of the pyoverdine biosynthetic gene *pvdA* is fused to the red fluorescent gene *mcherry*; and PAO1*pchEF*:*mcherry*, in which the promoter of the pyochelin biosynthetic genes *pchE* and *pchF* is fused to *mcherry*.

To simultaneously track the expression of two genes within an individual, we constructed double gene expression reporters. We constructed three different reporter strains involving the following pairs of genes: (1) PAO1*pvdA*::*mcherry–pchEF*::*egfp*, (2) PAO1*pchEF*::*mcherry–rpsL*::*egfp*, (3) PAO1*pvdA*::*mcherry–rpsL*::*egfp*. The genetic scaffold of the double reporter strain (1) is based on the construct proposed by Minoia et al. [37], where the two promoter fusions are located on the opposite strands, to minimize promoter interference (Fig. S1). For the double reporter strains (2) and (3), we developped an improved scaffold, where the promoter fusions are sequentially arranged on the leading strand (Fig. S2). Promoter interference was eliminated through the insertion of multiple terminator sites between the two promoter fusions. The bacterial strains, plasmids, and primers used in this study are listed in the supplementary Tables S2-S4, respectively.

### Growth conditions

Prior to experiments, overnight cultures were grown in 8ml Lysogeny broth (LB) in 50ml tubes, incubated at 37°C, 220 rpm for approximately 18 hours. Cells were harvested by centrifugation (8000 rpm for 2 minutes), subsequently washed in 0.8% saline and adjusted to OD600=1 (optical density at 600nm). For all batch culture and microscopy experiments, the harvested cells were grown in CAA medium (5g casamino acids, 1.18g K2HPO4*3H2O, 0.25g MgSO4*7H2O, per liter), buffered at physiological pH by the addition of 25mM HEPES. To induce iron limitation, we supplemented CAA with one of two iron chelator: the synthetic 2-2’-bipyridyl or the natural human apo-transferrin. While both iron chelators bind the extracellular iron, bipyridyl is also cell-permeable. The concentration ranges used were 50-300 μM for bipyridyl and 6.25-100 μg/ml for apo-transferrin. In the case of apo-transferrin, we also supplemented CAA medium with 20mM NaHCO3, an essential co-factor [38]. We further created an iron-replete condition by supplementing CAA with 100μM FeCl3. All chemicals were purchased from Sigma Aldrich (Buchs SG, Switzerland).

### Population-level growth and siderophore gene expression

We first quantified growth and siderophore gene expression of *P. aeruginosa* at the population level, by growing bacteria in CAA medium across a gradient of iron limitation (seven different conditions each for the experiments with either bipyridyl or apo-transferrin). We used the wildtype PAO1, and the single gene expression reporters PAO1*pvdA*:*mcherry* and PAO1*pchEF*:*mcherry*. Experiments were carried out in 200μl of medium distributed on a 96-well plate. Bacteria were inoculated at a starting density of OD600 = 0.0001, and incubated at 37°C for 24 hours in a multimode plate reader (Tecan, Männedorf, Switzerland). Cultures were shaken every 15 minutes for 15 seconds prior to measuring growth (OD600) and mCherry fluorescence (excitation:582nm/emission:620nm). We applied blank subtractions and standardized the relative fluorescence units (RFU) by OD600 (RFU/OD600). Each experiment featured four replicates per growth condition and was repeated twice.

We found that the growth integral (area under the curve) over 24 hours did not significantly differ between the wildtype PAO1 and the two single gene reporter strains (for bipyridyl: F_5,162_=0.0158, p=0.9843; for transferrin: F_5,165_=0.1357, p=0.8732). We therefore combined data from all three strains for growth analyses (Figures 1+S3).

**Fig 1.**
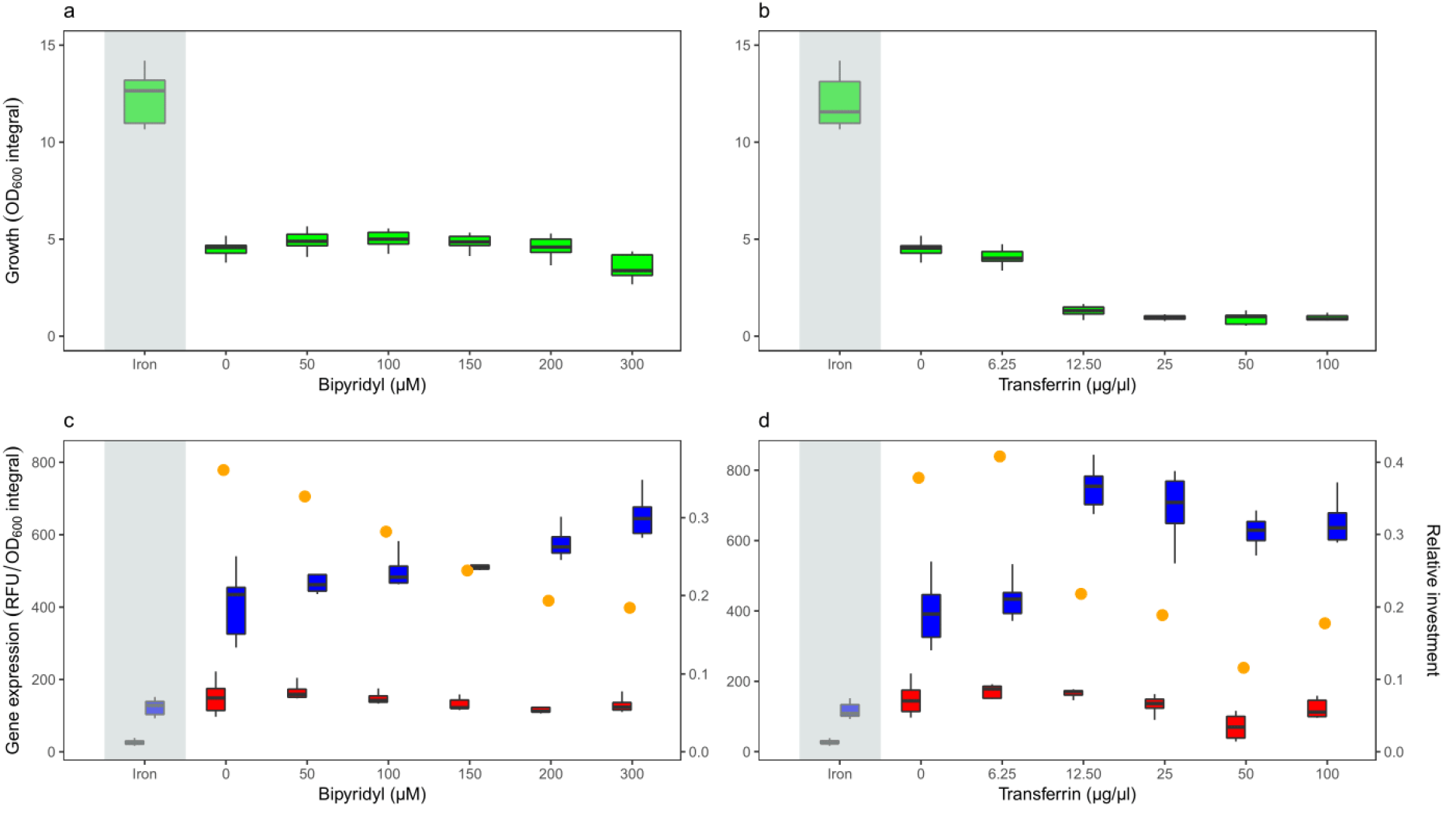
Higher levels of iron limitation reduce population growth and induce a gradual shift from pyochelin to pyoverdine gene expression in *P. aeruginosa*. **(a+b)** Growth measured as integral (area under the OD600 curve) over 24 hours batch culture experiments in iron-replete CAA medium (100μM FeCl3, grey shaded area) and CAA media with increasing concentrations of the iron chelators bipyridyl **(a)** and human apo-transferrin **(b)**. Higher iron chelator concentrations in the medium significantly reduced bacterial growth (see Table S1 for statistics). **(c+d)** Siderophore reporter gene expression (red: pyochelin PAO1*pchEF*:*mcherry*; blue: pyoverdine PAO1*pvdA*: *mcherry*) measured as normalized integral (area under the fluorescence signal curve divided by the OD600 integral) over 24 hours batch culture experiments. Gene expression was measured in iron-replete CAA medium (100μM FeCl3, grey shaded area) and CAA media with increasing concentrations of the iron chelators bipyridyl **(c)** and human apo-transferrin **(d)**. Relative gene expression was measured as the ratio of normalized pyochelin to pyoverdine expression (yellow dots). Boxplots represent the median with 25^th^ and 75^th^ percentiles, and whiskers show the 1.5 interquartile range.

### Siderophore gene expression at the single-cell level

We grew bacteria in batch cultures across a range of iron limitations in CAA (see above) in 1.5ml volumes distributed on 24-well plates, using a starting inoculum of OD600=0.0001. The cultures were incubated under shaken condition (170 rpm) for 24 hours at 37°C. From these growing cultures, we took small aliquots of 2μl every three hours (upto 24 hours), which we put on a microscopy slide for the quantification of single-cell gene expression using fluorescence microscopy. Because of the heavy workload associated with our 3-hours imaging interval across 24 hours, we split each experiment into two time blocks. We set up a first plate with experimental conditions from overnight LB cultures early in the morning of day 1 and measured gene expression of cells at times 3^rd^, 6^th^ and 9^th^ hour on day 1 and at the 21^st^ and 24^th^ hour on day 2. In addition, we transferred the overnight culture to fresh LB medium in the morning of day 1, to keep a growing bacterial population. This culture was then used to set up a second plate with experimental conditions in the evening of day 1. We used this second plate to measure gene expression of cells at the 12^th^,15^th^,18^th,^ and 21^st^ hours on day 2.

We performed our single-cell experiments with a subset of growth conditions used for our population-level experiments. Specifically, we used CAA medium supplemented with 100μM FeCl3 as the iron-replete condition, and gradually increased iron limitation by either adding no iron, 100μM or 300μM of bipyridyl. We further repeated the experiments by adding either 6.25, 25 or 100μg/ml of apo-transferrin.

We had two main experimental blocks. In block 1, we examined pyochelin and pyoverdine gene expression (separately for bipyridyl and apo-transferrin). Each replicate within block 1 featured the following four strains: (1) PAO1*pvdA*:*mcherry* single reporter; (2) PAO1*pchEF*:*mcherry* single reporter; (3) PAO1*pvdA*::*mcherry–pchEF*::*egfp* double reporter; and (4) the wildtype PAO1. In block 2, we compared siderophore with house-keeping gene *rpsL* expression. Each replicate within block 2 featured the following four strains: (1) PAO1*pvdA::mcherry–pchEF::egfp;* (2) PAO1*pchEF*::*mcherry–rpsl*::*egfp;* (3) PAO1*pvdA*::*mcherry–rpsl*::*egfp;* and (4) the wildtype PAO1. Each experimental block featured two full replicates including all conditions and time points. Block 1 was repeated another two times including the early time points only to compensate for the fact that cell number was low during the early growth phase.

### Preparation of microscope slides

For microscopy, we adapted a method previously described by de Jong et al. [39] and Weigert and Kümmerli [40]. Standard microscope slides (76mm x 26mm) were sterilized with 70% ethanol. We used ‘Gene Frames’ (Thermo Fisher Scientific, Vernier, Switzerland) to prepare agarose pads on which bacteria were seeded. Each frame features a single chamber (17 mm x 28 mm) of 25 mm thickness. The frames are coated with adhesives on both sides so that they stick to the microscope slide and the coverslip. The sealed chamber is airproof, which prevents pad deformation and evaporation during experimentation.

To prepare agarose pads, we heated 50 ml of 25mM HEPES buffer with agarose (1%) in a microwave. The agarose-buffer solution was cooled to approximately 50°C. We pipetted 700 μl of the solution into the gene frame and immediately covered it with a sterile coverslip. The coverslip was gently pressed to let superfluous medium escape. Slides were stored overnight at 4°C to ensure pad solidification. We removed the coverslip (by carefully sliding it sideways) and divided the agarose pad into six smaller pads of roughly equal size with a sterile scalpel. We introduced channels between pads, which served as oxygen reservoirs. Every pad received 2μl of a growing bacterial culture (of a specific time point, strain and experimental condition) extracted from the 24 well plates. We diluted cells in 0.8% saline to get an optimal number of individually discernible cells. The dilutions ranged from 1:1 at 3 hours to 1:100 at 24 hours. Upon the addition of bacteria, we let the agarose pads air-dry for 2 minutes, and then sealed them with a new sterile coverslip.

### Microscope set-up and imaging

We immediately imaged the bacteria on the pads at the Center for Microscopy and Image Analysis of the University of Zurich (ZMB) using an inverted widefield Olympus ScanR HCS microscope featuring the OLYMPUS cellSens Dimensions software. Images were captured with a PLAPON 60x phase oil immersion objective (NA=1.42, WD=0.15mm) and a Hamamatsu ORCA_FLASH 4.0V2, high sensitive digital monochrome scientific cooled sCMOS camera (resolution: 2048×2048 pixels, 16-bit). For fluorescence imaging, we used a fast emission filter wheel, featuring a FITC SEM filter for eGFP (excitation=470±24 nm, emission=515±30nm, DM=485) and a TRITC SEM filter for mCherry (excitation=550±15, emission=595±40, DM=558). We imaged at least six fields of view per pad, with each pad representing a specific combination of bacterial strain, experimental condition and time point.

### Image processing

For image processing, we established a semi-automated workflow [40]. The workflow starts with the machine learning, supervised object classification, and segmentation tool ILASTIK [41]. It features a self-learning algorithm that classifies objects (cells in our case) from the background using phase-contrast images. We used around 20 representative phase-contrast images from our experiments to train ILASTIK. After training, we supervised the results, marked errors and, re-initiated the next training round, until segmentation was optimized and nearly error-free. We then used the trained algorithm to segment all our images in a fully automated process [40]. Subsequently, we processed the segmented phase-contrast images with the open-source software FIJI [42], where regions of interest (ROI) were defined, capturing cells. We extracted information on cell size and fluorescence intensity for every single cell. To exclude segmented artifacts like cell debris, we set a cut-off area for ROIs, below which segmented objects were excluded.

We applied three different correction steps to our fluorescence images. The three steps were carried out independently for the green (eGFP) and red (mcherry) fluorescence channels. First, we corrected for agarose pad autofluorescence. For each agarose pad, we imaged at least 4 empty random positions without bacteria and averaged the grey values across pixels in the respective fluorescent channel. This average grey value was then subtracted from the fluorescence images containing cells. Then we overlaid the ROIs obtained from the phase-contrast images with the corresponding fluorescence images. Second, we corrected for intensity difference across the fields of view caused by microscope vignetting. To achieve this, we quantified the average grey value of the area outside the ROIs of each image containing cells, and subtracted that average grey value from all ROIs in that particular image. At the end of this step, we obtained background-corrected fluorescence values for each individual cell (i.e. ROI). We used the Integrated Density (IntDen) values, which is the mean grey value multiplied by the area of the cell, for further analysis. In the last step, the IntDen values were corrected for the auto-fluorescence of bacterial cells. Specifically, we measured the fluorescence of PAO1 wildtype cells that did not have a fluorescence reporter. Since the wildtype strain always ran in parallel with the reporter strains, we obtained measures of auto-fluorescence as IntDen values for a large number of wildtype cells for each time point and experimental condition. We log-transformed the IntDen values of wildtype cells and calculated the log(IntDen) median fluorescence value across all wildtype cells per time point and condition. We then subtracted these log(IntDen) median values from all individual cells of the respective time point and condition. A consequence of this correction procedure is that the log(IntDen) median value for the wildtype equals zero for all time points and conditions, and log(IntDen) values > 0 indicate gene of interest is expressed. Using this procedure, we quantified gene expression of 327 113 cells.

### Data stitching and statistical analysis

Each experimental block was repeated two to four times. While the results between repeats were reproducible in terms of relative gene expression patterns, the absolute level of gene expression naturally varied between repeats. For quantitative data analysis, we aimed to fuse the data sets from the individual repeats. This required a data normalization step. First, we calculated the mean gene expression *x_r_* for every condition, time point, and strain within each repeat. Next, we calculated the global mean 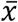 across repeats for the condition, time point and strain in question. Then, we calculated the difference between the two means 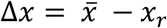. Finally, we subtracted Δ*x* from every single cell from the corresponding condition, time point and strain.

We used general linear models for statistical data analysis in R 3.4.2. For our population-level growth and gene expression analysis, we used analysis of variance (ANOVA) models with strain type as fixed factor and iron chelator concentration as co-variate. We conducted separate analyses for the two iron chelators bipyridyl and apo-transferrin. To test whether pyochelin and pyoverdine gene expression varies across time categories, we conducted one-way ANOVAs for each media condition separately, subsequently, we performed a posthoc Tukey honest significant difference (HSD) test for pairwise comparison between time categories. Next, we compared the standard deviation in gene expression across cells, as a measure of heterogeneity, across time and media conditions. We used ANOVA models, where we fitted media condition as a fixed factor and time as a covariate. Finally, we calculated Pearson’s correlation coefficient *r* to relate the expression of two genes across cells for any given condition tested. We then extracted all *r*-values and used ANOVAs to test whether the strength of correlations varies as a function of media condition (fixed factor) and time (co-variate). Finally, we conducted paired *t*-tests to compare whether correlation coefficients differ between the expression of the two siderophore genes (*pchEF* vs. *pvdA*), and the expression of a siderophore gene (*pchEF* or *pvdA*) and the housekeeping gene rpsL. We log-transformed all single-cell gene expression values prior to analysis.

## RESULTS

### Population-level growth and siderophore gene expression

We first quantified growth and siderophore gene expression of *P. aeruginosa* at the population level, across a range of CAA media compositions, differing in their level of iron limitation. As expected, we found that iron supplementation increased population growth, while the addition of iron chelators reduced growth relative to the plain CAA medium (Fig. 1a+b, Table S1). However, the two iron chelators used (bipyridyl and human apo-transferrin) varied in their effect on growth. The addition of bipyridyl predominantly reduced the lag-phase of cultures (Fig S3, Table S1, F_1,238_=144.9, p<0.0001), but had only a mild negative effect on the growth integral (F_1,238_=18.08, p<0.0001) and no significant effect on the maximal growth rate (Fig S3, Table S1, F_1,238_=1.65, p=0.2002). Conversely, the addition of transferrin significantly reduced all three growth parameters (Fig S3, Table S1, lag-phase: F_1,298_=252.4, p<0.0001; growth integral: F_1,298_=474.6, p<0.0001; maximal growth rate: F_1,298_=121.6, p<0.0001).

With regard to gene expression, we found very low fluorescence signals for pyochelin (PAO1*pchEF*:*mcherry*) and pyoverdine (PAO1*pvdA*:*mcherry*) in iron-supplemented medium (Fig. 1c+d), confirming that the investment in the two siderophores is reduced to baseline levels [30, 43]. Conversely, pyochelin and pyoverdine genes were expressed in plain CAA medium and CAA supplemented with iron chelators. While pyoverdine gene expression significantly increased with higher chelator concentrations (for bipyridyl: F_1,79_=108.3, p<0.0001; for transferrin: F_1,78_=64.52, p<0.0001), pyochelin gene expression significantly decreased (for bipyridyl: F_1,77_=8.77, p=0.0041; for transferrin: F_1,78_=17.96, p<0.0001). Consequently, the ratio of pyochelin-to-pyoverdine gene expression significantly changed in favor of pyoverdine with higher levels of iron limitation (for bipyridyl: F_1,4_=162.4, p=0.0002; for transferrin: F_1,4_=12.05, p=0.0255). Temporal dynamics of population-level gene expression confirmed these patterns (Fig. 2): pyoverdine gene expression increased monotonously over time, while pyochelin gene expression plateaued or even declined over time with high chelator concentrations. These population level gene expression patterns confirm previous findings based on measurements of actual siderophore production [26], showing that bacteria gradually shift from pyochelin to pyoverdine production when moving from mild to more severe iron limitation.

**Fig 2.**
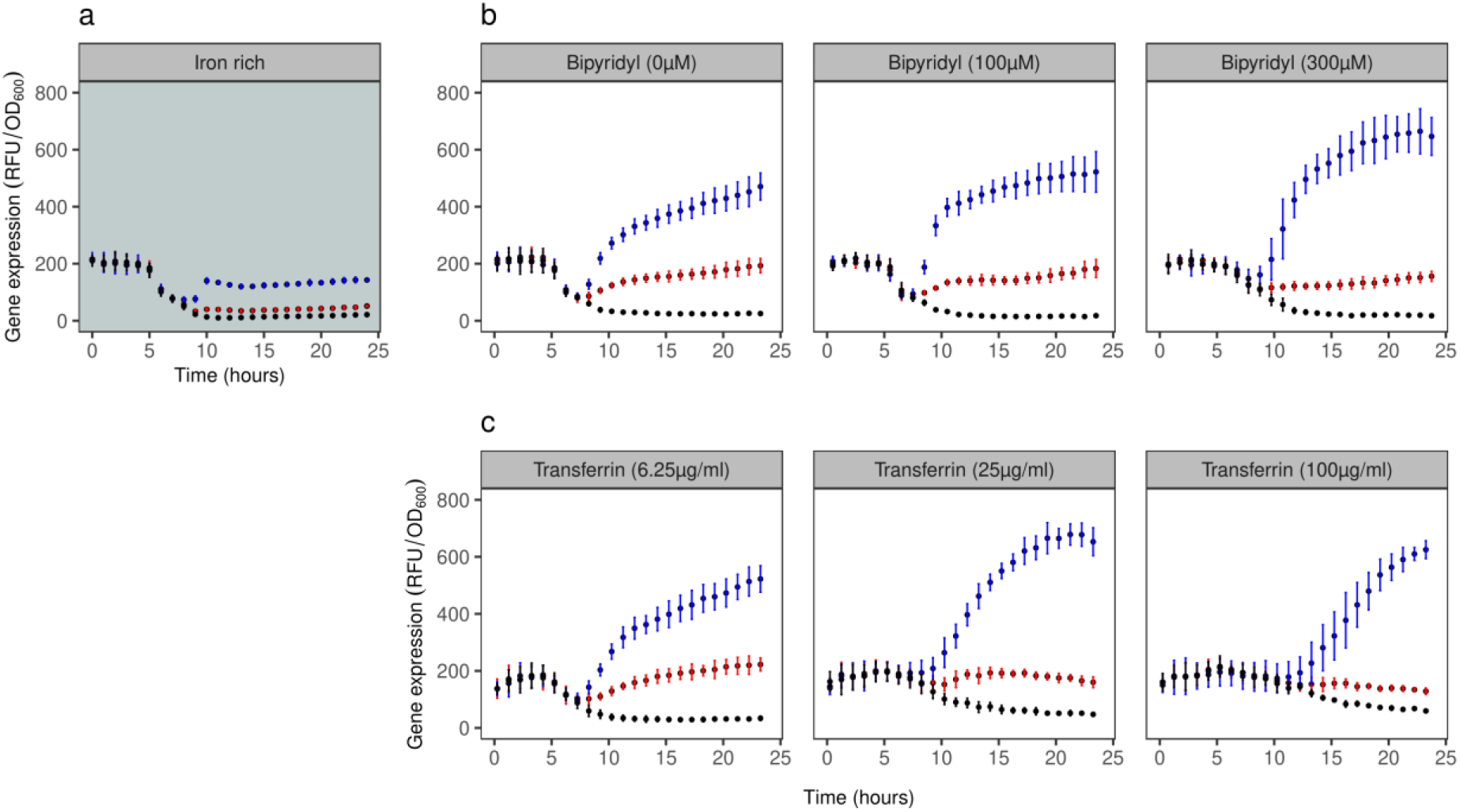
Temporal dynamics of siderophore reporter gene expression at the population level. The panels show reporter gene expression for pyochelin (red: PAO1*pchEF*:*mcherry*), pyoverdine (blue: PAO1*pvdA*:*mcherry*), and the wildtype control strain (black: PAO1 without reporter fusion) across a range of CAA media differing in their level of iron limitations. Values represent normalized gene expression (fluorescence signal divided by OD600) in: **(a)** iron-replete CAA medium (100 μM FeCl3); **(b)** CAA media with increasing concentrations of the iron chelator bipyridyl; **(c)** CAA media with increasing concentrations of the iron chelator apo-transferrin. Values and error bars represent the mean and standard deviation across 8 replicates (2 experiments with 4 replicates each), respectively. The inflation of normalized gene expression in the early hours of the growth can be attributed to divisions by very low OD600 values (close to zero) in this growth phase. Gene expression patterns started to segregate from the control PAO1 strain upon entering the exponential growth phase.

### Single-cell level siderophore gene expression across time and iron limitations

We used the double reporter PAO1*pvdA*::*mcherry–pchEF*::*egfp* to quantify pyochelin and pyoverdine gene expression of individual cells over a 24 hours growth cycle in batch cultures, across seven different variants of the CAA medium differing in their level of iron availability. In the following sub-sections, we will dissect the complex expression patterns by first focusing on the iron-supplemented treatment, and then turning to iron-limited media supplemented with either bipyridyl or apo-transferrin. The single-cell data is shown in three different ways: as dot plots (Fig. 3, where each dot represents an individual cell), as density plots (Fig. S4), and as box plots (Fig. S5 for statistical comparisons). Note that we repeated all experiments with the single gene reporters PAO1*pvdA*:*mcherry* and PAO1*pchEF*:*mcherry* as controls, which run in parallel with our double reporter. The results from the single (Fig. S6) and double reporters (Fig. S5) are highly congruent, demonstrating that our findings are neither influenced by the construct type nor the fluorophore (mCherry vs. eGFP).

**Fig 3.**
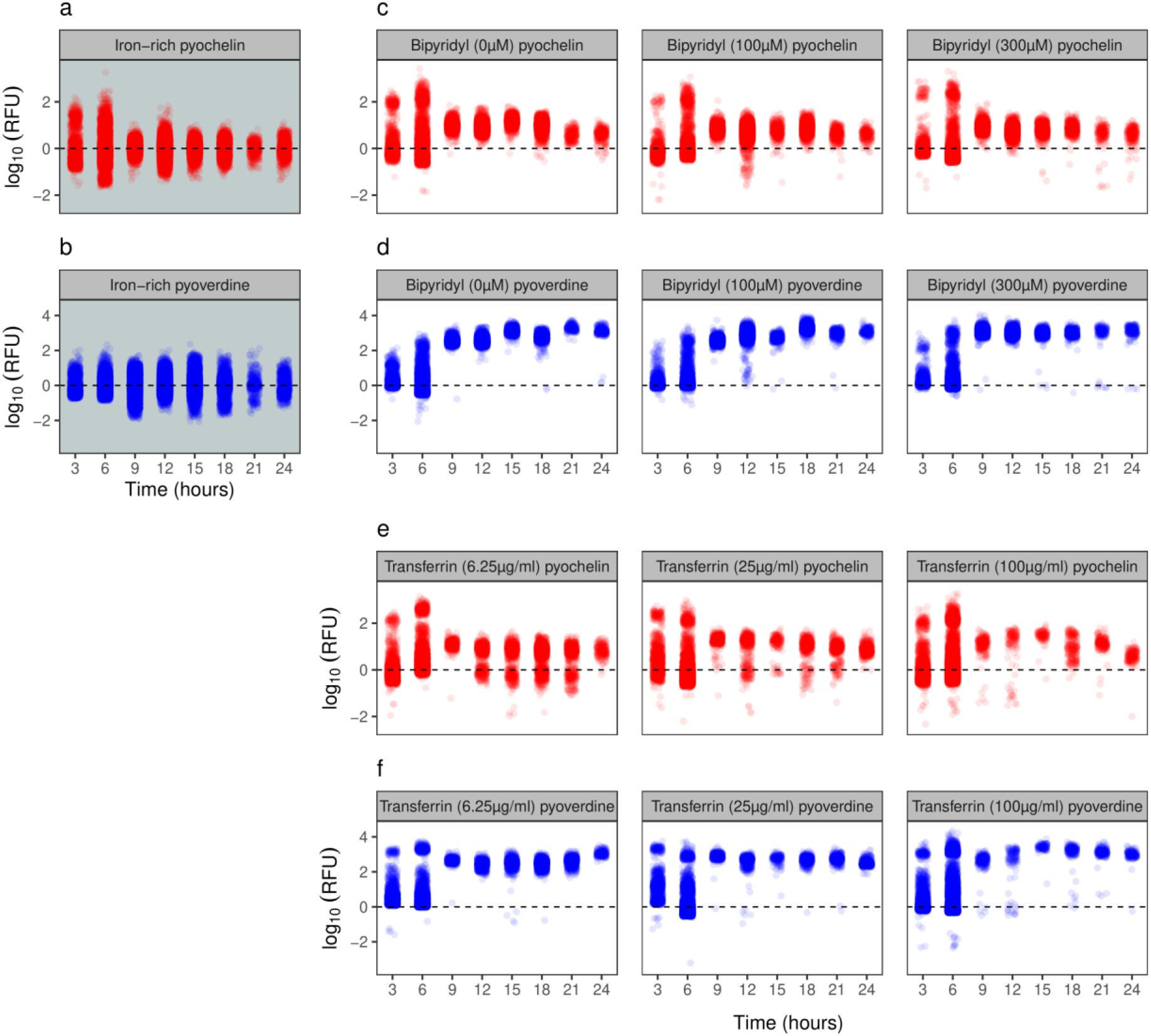
Single-cell siderophore gene expression patterns across time and media. The expression of pyochelin (red: *pchEF*) and pyoverdine (blue: *pvdA*) synthesis genes measured with the double reporter PAO1*pvdA*::*mcherry–pchEF*::*egfp* across a range of CAA media differing in their levels of iron limitations. Each dot represents an individual cell with gene expression shown as log-transformed fluorescence values. (**a**+**b**) Iron-replete CAA medium (100 μM FeCl3); (**c**+**d**) CAA media with increasing concentration of the iron chelator bipyridyl; (**e**+**f**) CAA media with increasing concentration of the iron chelator apo-transferrin.

#### Siderophore gene expression patterns in iron-rich medium

Consistent with previous batch-cultures studies [19, 23], we found that the average *pchEF* and *pvdA* gene expression did not differ from background levels in iron-rich CAA medium, regardless of the time points analyzed (Fig. S5a+b). However, gene expression varied widely between cells (Fig. 3a+b). For pyochelin, we observed bi-modal gene expression up to 6 hours, where a fraction of cells (16 %) showed relatively high *pchEF* expression activity (log(fluo) ≥ 1), whereas the remaining cells were primarily in the off stage (Fig. S4a). At later time points, the bimodality disappeared. When comparing standard deviations as a proxy for gene expression heterogeneity (Fig. 4a), we observed that *pchEF* expression heterogeneity peaked early on during the experiment (3-6 hours) and then declined afterwards. For pyoverdine, there was no bimodality in gene expression (Fig. S4b), but relatively high and time-consistent heterogeneity in *pvdA* gene expression across cells (Fig. 4a).

**Fig 4.**
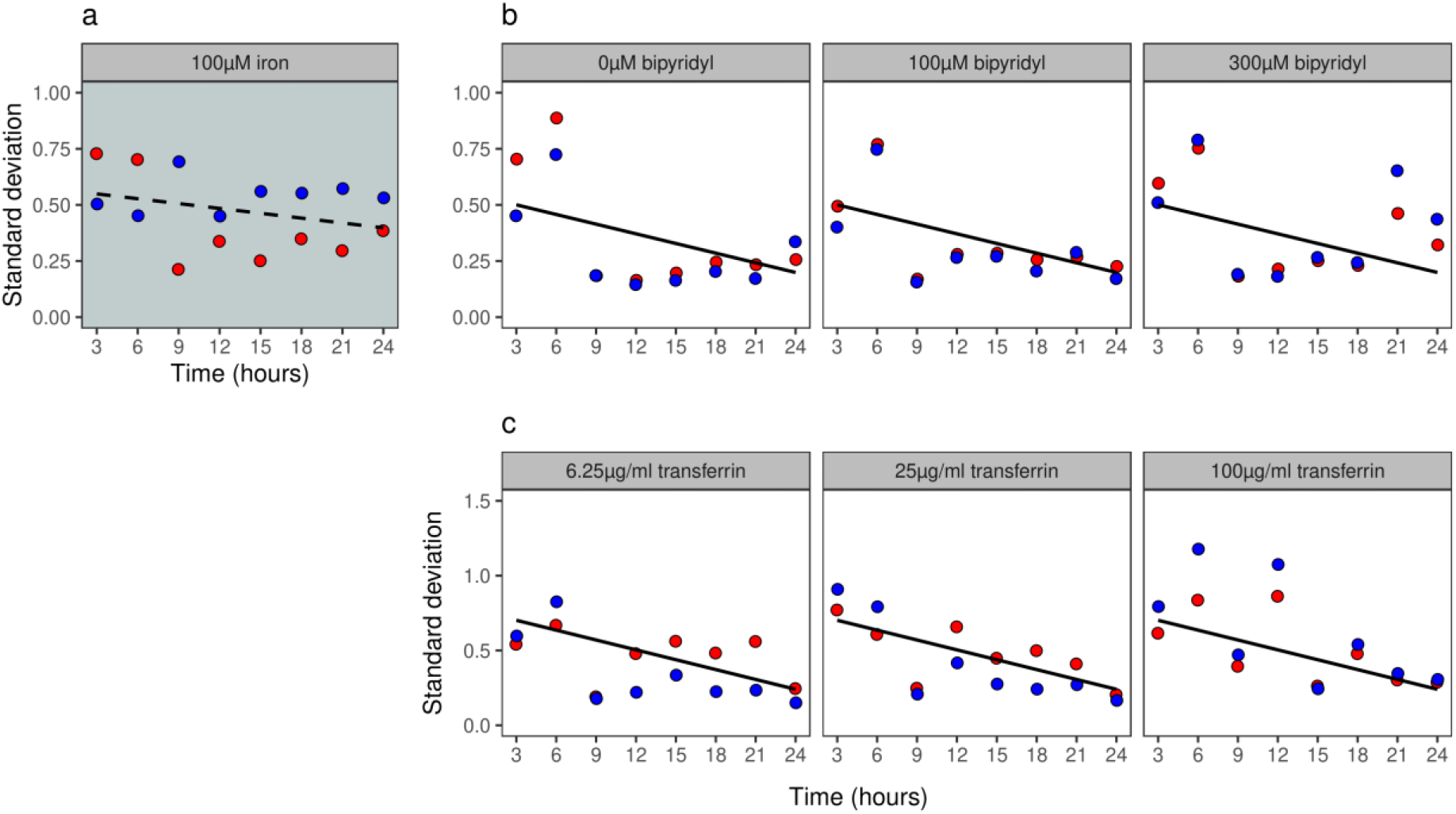
Heterogeneity in siderophore gene expression declines over time. Temporal heterogeneity in siderophore gene expression for pyochelin (red: *pchEF*) and pyoverdine (blue: *pvdA*), measured with the double reporter strain PAO1*pvdA*::*mcherry*–*pchEF*::*egfp* across a range of CAA media differing in their levels of iron limitations. Heterogeneity in gene expression is shown as the standard deviation across log-transformed fluorescence values of all cells. **(a)** Iron-replete CAA medium (100 μM FeCl3); **(b)** CAA media with increasing concentration of the iron chelator bipyridyl; **(c)** CAA media with increasing concentration of the iron chelator apo-transferrin. Solid black trendlines indicate significant declines in heterogeneity across time.

#### Siderophore gene expression patterns in media supplemented with bipyridyl

Compatible with our population-level analysis (Fig. 2), we found that *pchEF* gene expression peaked at intermediate time points (Fig. S5c), whereas *pvdA* expression was consistently high until the end of the experiment (Fig. S5d). Again, we observed that gene expression varied widely between cells (Fig. 3c+d). For pyochelin, we consistently found bi-modal *pchEF* expression during the first six hours of the experiment, regardless of how much bipyridyl was added (Fig. S4c). Under all conditions, there was a high and a low proportion of cells investing marginally and highly in pyochelin gene expression, respectively. At later time points, gene expression levels across cells converged to intermediate levels (Fig. 3c). For pyoverdine, the pattern was different in the sense that we observed a gradual induction of gene expression from an off-low to an all-on stage (Fig. 3d+S4d). We found that the heterogeneity in *pchEF* and *pvdA* expression significantly decreased over time (F_1,43_ =12.90, p=0.0008), with the decrease being similar for both genes (F_1,43_=0.13, p=0.7207) and across media (F_1,43_ =0.61, p=0.5500; Fig. 4b).

#### Siderophore gene expression patterns in media supplemented with apo-transferrin

The single-cell gene expression patterns in CAA media supplemented with apo-transferrin mirrored at large the patterns observed in media containing bipyridyl: (i) *pchEF* gene expression peaked at intermediate time points (Fig. S5e), whereas *pvdA* expression was highest at the end of the experiment (Fig. S5f); (ii) heterogeneity in gene expression across cells was high, especially during the first six hours of the experiment (Fig. 3e+f, Fig. S4e+f); and (iii) heterogeneity in *pchEF* and *pvdA* expression significantly decreased over time (F_1,43_=26.93, p<0.0001), similarly for both genes (F_1,43_=0.19, p=0.6625) and across media (F_1,48_=2.59, p=0.0865; Fig. 4c). However, there were also a few notable differences. First, bimodal gene expression during the first six hours of the experiment did not only occur for *pchEF* (Fig. 3e+S4e), but also for *pvdA* (Fig. 3f+S4f). Moreover, the bimodality in *pchEF* expression persisted at many of the later time points, especially under conditions with low apo-transferrin concentration (Fig. 3e+S4e). Taken together, the single-cell analysis reveals a funneling pattern in the expression of siderophore synthesis genes in all iron-limited media, from high between-cell heterogeneity (including bimodality) during the early time points to greatly aligned homogenous expression patterns during later time points. Moreover, there was a temporal shift from pyochelin to pyoverdine gene expression under iron-limited conditions.

### Positive correlations between pyochelin and pyoverdine gene expression across cells

We then tested for correlations in the expression of pyochelin and pyoverdine synthesis genes across cells. Positive correlations would indicate that individuals in a clonal population segregate along a continuum from low to high siderophore producers, while negative correlations would suggest that cells specialize in either pyochelin or pyoverdine production. We observed positive correlations between pyochelin and pyoverdine gene expression for all time points and media conditions (Fig. 5a+b, Fig. S7), refuting the specialization but supporting the continuum hypothesis.

**Fig 5.**
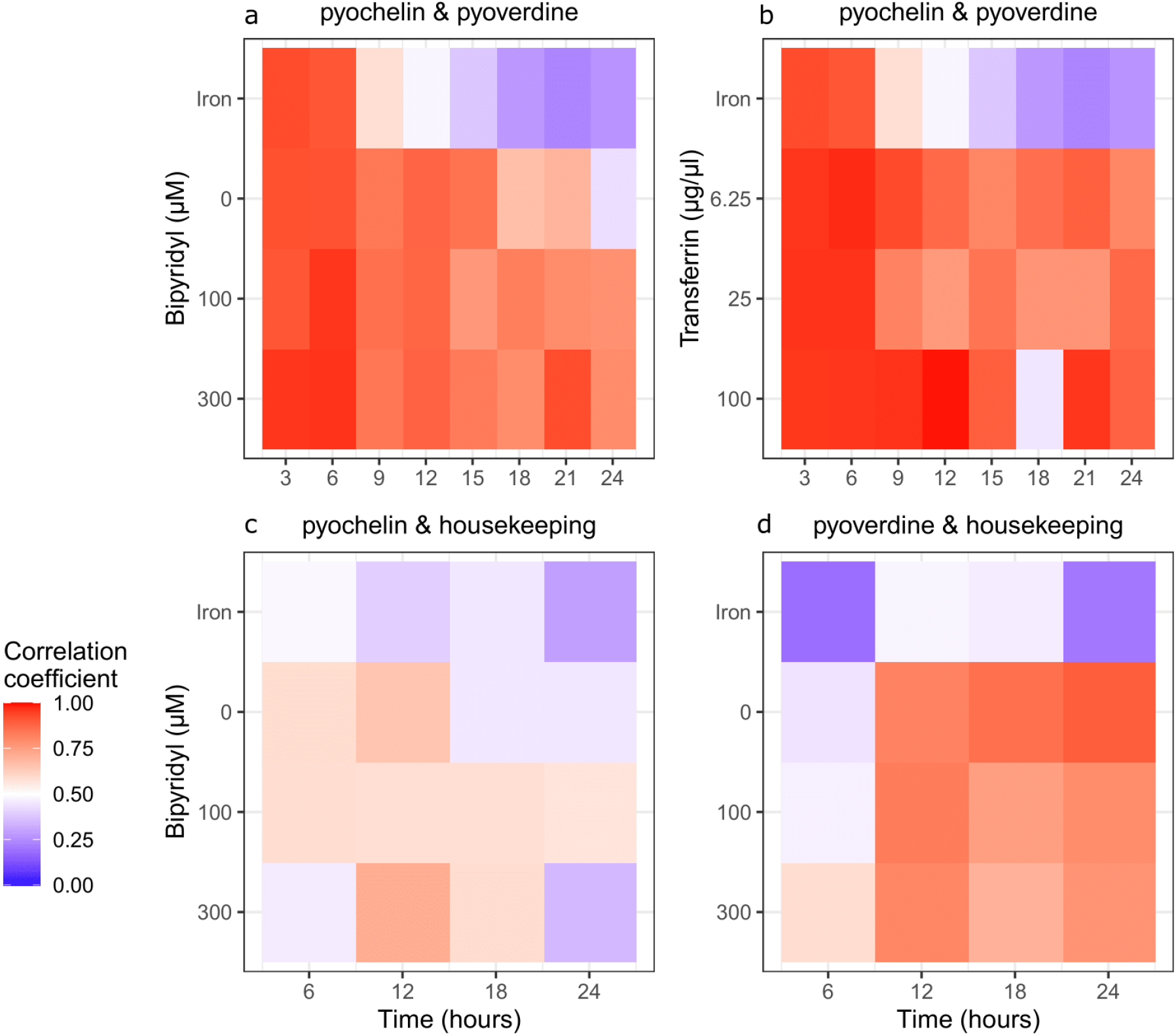
Correlations between pyochelin, pyoverdine and housekeeping gene expression among clonal cells across time and media. Correlation coefficients are shown as heatmaps, and indicate the strength of associations in the expression of two genes across individual cells within a population. **(a + b)** Correlations between pyochelin (*pchEF*) and pyoverdine (*pvdA*) gene expression, measured with the double reporter strain PAO1*pvdA*::*mcherry–pchEF*::*egfp* in iron-supplemented CAA medium and in CAA media supplemented with either increasing concentrations of the iron chelators bipiridyl or apo-transferrin. **(c)** Correlations between pyochelin *pchEF* and housekeeping *rpsL* gene expression, measured with the double reporter PAO1*pchEF*::*mcherry*–*rpsL*::*egfp* in iron-supplemented CAA medium and in CAA media supplemented with increasing concentrations of the iron chelator bipyridyl. **(d)** Correlations between pyoverdine *pvdA* and housekeeping *rpsL* gene expression measured with the double reporter PAO1*pvdA*::*mcherry–rpsL*::*egfp* in iron-supplemented CAA medium and in CAA media supplemented with increasing concentrations of the iron chelator bipiridyl.

Under the iron-supplemented condition, the positive correlations were initially strong, but then significantly declined over time (F_1,6_=46.57, p=0.0005). In CAA medium with bipyridyl, the correlation coefficients also significantly declined over time (ANCOVA: F_1,18_=34.66, p<0.0001), with the decline being steeper under less stringent iron limitation (significant interaction between time and media: F_2,18_=5.423, p=0.0143). In CAA medium supplemented with apo-transferrin, the correlation coefficient also significantly declined over time (F_1,20_=5.061, p=0.0359), and similarly so for all conditions (F_2,20_=0.326, p=0.7254).

To further explore the continuum hypothesis, we examined whether the observed positive correlations arise because cells in clonal populations differ in their overall metabolic state, and whether the more active cells are the ones that show higher investments into siderophores. To test this idea, we simultaneously quantified the expression of one of the siderophore genes (*pchEF* or *pvdA*) together with the *rpsL* housekeeping gene within the same cell. In this context, we use *rpsL* housekeeping gene expression as a proxy for the overall metabolic activity of a cell. We observed positive correlations between *rpsL* and both *pchEF* (Fig. 5c, Fig. S8a) and *pvdA* (Fig. 5d, Fig. S8b) expression, for all time points and media conditions. These results strongly suggest that the overall metabolic state of a cell dictates its siderophore investment levels. However, we also observed that the correlation coefficients were significantly weaker for the *pchEF*-*rpsL* than for the *pchEF*-*pvdA* gene pair (paired t-test: *t*11=6.70, p<0.0001), while they were not different between *pvdA-rpsL* and *pchEF-pvdA* gene pairs (*t*11=1.14, p=0.2780). These findings indicate that investment into the larger pyoverdine depends more stringently on the metabolic state of the cell than the smaller pyochelin.

## DISCUSSION

The aim of our study was to obtain a detailed view on siderophore gene expression at the single cell level in growing populations of the bacterium *Pseudomonas aeruginosa*. This species produces two siderophores, pyochelin and pyoverdine, and the regulatory mechanisms governing the expression of these secondary metabolites are well known [22]. However, unknown is whether the seemingly fine-tuned regulation of siderophores, in response to iron availability and cell density, observed at the population level is driven by coordinated homogenous behaviours of the individual cells in the population. Our null-hypothesis was that there is no coordination such that the population-level response simply reflects the sum of its noisy individuals [17]. We could refute this hypothesis, as we observed that cells switched from initially heterogenous to highly homogenized gene expression patterns over the population growth cycle. One of our alternative hypothesis was that coordination could promote specialization, whereby subpopulations of cells either invest in pyoverdine or pyochelin [33]. We could also refute this hypothesis, as we found no evidence for negative correlations in the expression of the two traits across cells. Instead, our data strongly support that cells switch from an initially uncoordinated phase to highly coordinated, temporally stratified gene expression patterns in growing populations. The observed patterns are consistent across different iron-limited environments and are reminiscent of chronobiological (cyclical physiological) processes known from multi-cellular organisms [44, 45].

Our results suggest that the chronobiological cycle of siderophore gene expression entails three different phases, covering the time spans from low to high population density. We here propose that the three phases (I to III) are steered by the various interconnected regulatory mechanisms governing siderophore synthesis (Fig. 6). Phase I comprises the first six hours of our experiment, during which both pyochelin and pyoverdine gene expression are highly heterogenous across cells. Heterogeneity is often characterized by bimodal gene expression, where the majority of cells remains in the off-stage, while a minority of cells shows high (often overshooting for pyochelin) siderophore gene expression. We argue that this bimodality is driven by differences in the internal iron stocks of cells (Fig. 6a). Our cells are taken from iron-rich overnight LB cultures such that the acquired iron stocks should allow survival and growth in iron-limited media for a short time period [46]. Once the iron stocks are depleted, Fur-mediated repression is released and siderophore gene expression starts [20, 23–25]. Since each cell responds individually to its internal iron stocks, bimodality can most parsimoniously be explained by inter-individual variation in iron stocks. Phase II involves the timeframe from 6-15 hours, during which heterogeneity in siderophore gene expression is greatly reduced, and all cells switch to an on-state. Phase II coincides with the exponential growth phase of populations (Fig. S3). We argue that siderophore-mediated signaling of cells triggers the homogenization of gene expression (Fig. 6b). Signaling occurs when iron-loaded siderophores bind to membrane-bound cognate receptors [27, 28]. Signaling is likely stochastic at low population density and low siderophore concentrations as experienced in Phase I. Meanwhile, signaling is likely more deterministic in Phase II, where siderophore concentrations and cell density increases, and the signals can be shared more reliably between cells, leading to the homogenization of behaviors across cells [14]. Phase III involves the time points from 15 hours onwards, where cells continue growing at higher cell densities (Fig. S3). Here, we observed a shift in the relative gene expression from pyochelin to pyoverdine, while heterogeneity across cells remained low (Fig. 6c). While the low heterogeneity is likely the result of ongoing siderophore-signalling, we argue that the shift towards reduced pyochelin gene expression is governed by the hierarchical regulatory link between the siderophores, where high pyoverdine inhibits pyochelin synthesis [26, 29]. Although the exact mechanism of this inhibition is unknown, our data suggest that it occurs concomitantly in all cells. Taken together, the integration of three regulatory elements (Fur, siderophore-signalling and hierarchy) could guide the chronobiological cycle of siderophore gene expression in populations of *P. aeruginosa*, a pattern that is expected to be reinitiated in each new growth cycle.

**Fig 6.**
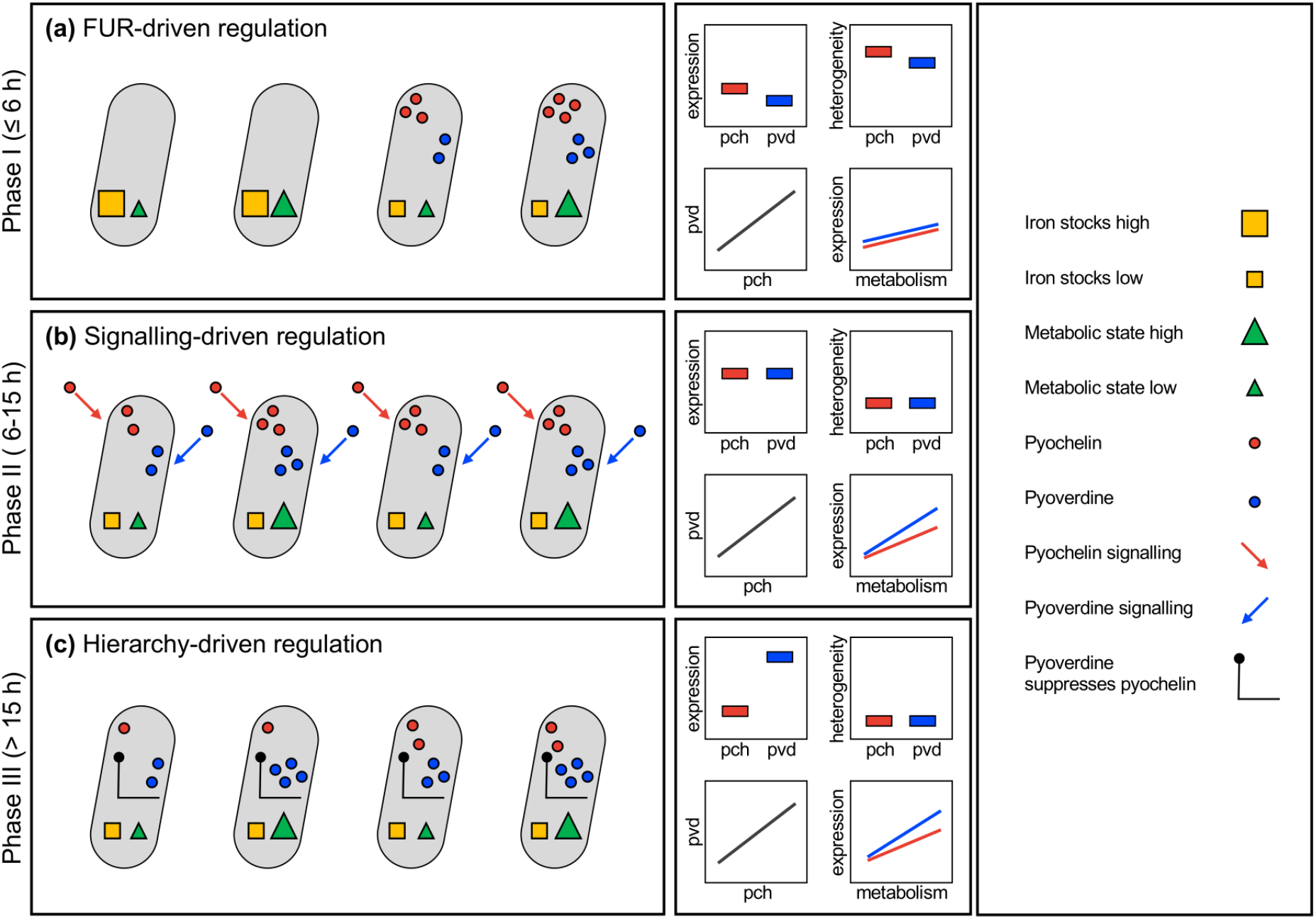
A three-phase model of how regulatory elements, internal iron stocks and the metabolic state of cells can explain decision-making processes and the chronobiological cycle of siderophore gene expression in growing populations of *P. aeruginosa*. **(a)** Phase I (up to 6 hours) is driven by Fur (ferric uptake regulator) and variation in internal iron-stocks across cells. Cells with low stocks start to express and produce pyochelin and pyoverdine regardless of their metabolic state. This induces high gene expression heterogeneity (including bimodality) across cells and strong positive correlations between pyochelin and pyoverdine expression. **(b)** Phase II (between 6 and 15 hours) is driven by signalling-mediated regulation and variation in the metabolic state. Siderophore-mediated signalling leads to homogenized and increased levels of gene expression. Residual variation across cells is largely explained by metabolic state differences. **(c)** Phase III (> 15 hours) is driven by hierarchy-driven regulation, inducing a shift from pyochelin to pyoverdine gene expression. Residual variation across cells is still explained by metabolic state differences, and more so for pyoverdine than for pyochelin.

Despite the fact that siderophore gene expression became homogenized across cells over time, there was still considerable inter-individual variation (Fig. 3). Our analyses suggest that this variation is largely explained by differences in the metabolic state of cells. Our reasoning is based on the observation that the expression of both siderophore genes (*pchEF* and *pvdA*) correlated positively with the *rpsL* housekeeping gene expression (Fig. 5). *RpsL* encodes the 30S ribosomal protein S12, which plays a major role in translational accuracy. While *rpsL* is essential and expressed by all cells, the expression magnitude can be taken as a proxy for metabolic activity [47]. Interestingly, we observed that the strength of the positive correlations between *rpsL* and siderophore genes varied across phases I – III and also differed between the siderophores (Fig. 5+6). In phase I, the association between *rpsL* and siderophore gene expression was relatively weak. This suggests that siderophore gene expression is predominantly triggered by Fur de-repression and not the metabolic state (Fig. 6a). In contrast, stronger correlations between metabolic activity and siderophore gene expression arose in phase II and III. Since these phases cover the exponential growth period during which cells are metabolically most active, our data suggest that cells reaching a higher metabolic state make more siderophores. Finally, we found that the expression of pyoverdine was more strongly correlated with metabolic activity than pyochelin expression (Fig. 5c+d). This weaker association could arise because the small pyochelin is cheaper to produce than the large pyoverdine, meaning that the production of pyochelin is less dependent on the metabolic state of a cell [26].

While the above sections offer mechanistic explanations for the observed siderophore gene expression patterns, we here aim to discuss possible adaptive (evolutionary) reasons for why bacteria behave the way they do. An obvious explanation is that the specific regulatory circuits have evolved to optimize the cost-to-benefit ratio of siderophore investment under iron-limited conditions [26, 31]. However, there seems to be more to it. For example, our data suggest that the role of pyochelin might have been underestimated in the past [29]. While typically considered as a secondary siderophore, we show that pyochelin gene expression occurs under all conditions and peaks early during the growth cycle, while pyoverdine gene expression kicks in later. One adaptive explanation could be that bacteria follow a two-step iron-scavenging strategy when colonizing a new patch. They first invest in the cheap pyochelin during the early colonization phase, where population density is low and many siderophores might get lost due to diffusion [5]. They then switch to the more expensive pyoverdine when reaching higher population densities, where the sharing of siderophores becomes more reliable and fewer molecules are lost. The ‘early-on’ scavenging role of pyochelin is further supported by our observation that there is always a fraction of cells in the population that have pyochelin expression switched on, even under iron rich conditions. This could point towards a bet-hedging strategy [18, 48, 49], where clonal populations maintain a fraction of cells in an pyochelin-on state, in order to be immediately able to react to an environmental change in iron limitation. This strategy could provide a competitive edge over a strategy, where all cells are in a siderophore-off stage, because it cuts short the initial sensing of iron limitation and the mounting siderophore synthesis from scratch [50].

As a cautionary note, it is important to consider that we solely looked at gene expression and not at siderophore production itselves. Post-transcriptional regulation and translational noise could weaken the assumed positive correlation between gene expression and siderophores synthesis. However, we have high confidence that such positive correlations are real, because our population-level gene expression data (Fig. 1) match actual siderophore production levels from our previous work [26, 51].

In summary, our single-cell gene expression study highlights that clonal bacterial population can coordinate their actions in ways that match decision-making processes observed in higher organisms [52–55]. Specifically, decision-making theory predicts that individuals can coordinate their actions when integrating information from their local neighbors. Moreover, the response precision can be increased when information is sampled across longer time periods or a broader range (e.g., across more individuals) [14, 56]. Siderophore signaling fulfils the function of information collection with the aggregate availability of siderophores at the local group level being the currency of information. It provides information on cell density and the siderophore investment of neighbors. The accuracy of information collection increases with cell density and could explain why the coordinated response only emerges after some time. Overall, our single-cell approach offers a deeper understanding on the coordination of bacterial social behaviors among cells in clonal groups. While we used siderophores as a model trait, we believe that our methods and concepts can also be applied to study coordination of any other bacterial social trait.

## Acknowledgments

We thank Priyanikha Jayakumar, Chiara Rezzoagli, Tobias Wechsler, Jos Kramer and Michael Weigert for help in the lab and with the data analysis, and the Center for Microscopy and Image Analysis for support. This project has received funding from the European Research Council (ERC) under the European Union’s Horizon 2020 research and innovation programme (grant agreement no. 681295).

## Author contributions

SM and RK designed the study. SM carried out all experiments and constructed some of the strains. SM and RK analyzed the data and wrote the paper.

## References

1. West SA, Diggle SP, Buckling A, Gardner A, Griffin AS. The social lives of microbes. Annu Rev Ecol Evol Syst. 2007;38:53–77.

2. Diggle SP, Griffin AS, Campell GS, West SA. Cooperation and conflict in quorum-sensing bacterial populations. Nature. 2007;450:411–414.

3. Ebrahimi A, Schwartzman J, Cordero OX. Cooperation and spatial self-organization determine rate and efficiency of particulate organic matter degradation in marine bacteria. Proc Natl Acad Sci USA. 2019;116:23309–23316.

4. Yan J, Monaco H, Xavier JB. The ultimate guide to bacterial swarming: an experimental model to study the evolution of cooperative behavior. Annual Reviews in Microbiology. 2019;73:293–312.

5. Kramer J, Özkaya Ö, Kümmerli R. Bacterial siderophores in community and host interactions. Nat Rev Microbiol. 2020;18:152–163.

6. Griffin A, West SA, Buckling A. Cooperation and competition in pathogenic bacteria. Nature. 2004;430:1024–1027.

7. Sandoz KM, Mitzimberg SM, Schuster M. Social cheating in *Pseudomonas aeruginosa* quorum sensing. Proc Natl Acad Sci USA. 2007;104:15876–15881.

8. Xavier JB, Kim W, Foster KR. A molecular mechanism that stabilizes cooperative secretions in *Pseudomonas aeruginosa*. Mol Microbiol. 2011;79:166–179.

9. Drescher K, Nadell CD, Stone HA, Wingreen NS, Bassler BL. Solutions to the public goods dilemma in bacterial biofilms. Curr Biol. 2014;24:50–55.

10. Nadal Jimenez P, Koch G, Thompson JA, Xavier KB, Cool RH, Quax WJ. The multiple signaling systems regulating virulence in *Pseudomonas aeruginosa*. Microbiol Mol Biol Rev. 2012;76:46–65.

11. Schuster M, Sexton DJ, Diggle SP, Greenberg EP. Acyl-homoserine lactone quorum sensing: from evolution to application. Annu Rev Microbiol. 2013;67:43–63.

12. Papenfort K, Bassler BL. Quorum sensing signal-response systems in Gram-negative bacteria. Nat Rev Microbiol. 2016;14:576–588.

13. Darch SE, West SA, Winzer K, Diggle SP. Density-dependent fitness benefits in quorum-sensing bacterial populations. Proc Natl Acad Sci USA. 2012;109:8259–8263.

14. Ross-Gillespie A, Kümmerli R. Collective decision-making in microbes. Front Microbiol. 2014;5:54.

15. Whiteley M, Diggle SP, Greenberg EP. Progress in and promise of bacterial quorum sensing research. Nature. 2017;551:313–320.

16. Avery A. Microbial cell individuality and the underlying sources of heterogeneity. Nat Rev Microbiol. 2006;4:577–587.

17. Eldar A, Elowitz MB. Functional roles for noise in genetic circuits. Nature. 2010;467:167–173.

18. Ackermann M. A functional perspective on phenotypic heterogeneity in microorganisms. Nat Rev Microbiol. 2015;13:497–508.

19. Visca P, Imperi F, Lamont IL. Pyoverdine siderophores: from biogenesis to biosignificance. Trends Microbiol. 2007;15:22–30.

20. Youard ZA, Wenner N, Reimmann C. Iron acquisition with the natural siderophore enantiomers pyochelin and enantio-pyochelin in *Pseudomonas* species. Biometals. 2011;24:513–522.

21. Schalk IJ, Cunrath O. An overview of the biological metal uptake pathways in *Pseudomonas aeruginosa*. Environ Microbiol. 2016;18:3227–3246.

22. Schalk IJ, Rigouin C, Godet J. An overview of siderophore biosynthesis among fluorescent Pseudomonads and new insights into their complex cellular organization. Environ Microbiol. 2020;22:1447–1466.

23. Ochsner UA, Vasil ML. Gene repression by the ferric uptake regulator in *Pseudomonas aeruginosa*: cycle selection of iron-regulated genes. Proc Natl Acad Sci USA. 1996;93:4409–4414.

24. Leoni L, Ciervo A, Orsi N, Visca P. Iron-regulated transcription of the *pvdA* gene in *Pseudomonas aeruginosa*: effect of Fur and PvdS on promoter activity. J Bacteriol. 1996;178:2299–2313.

25. Escolar L, Pérez-Martín J, de Lorenzo V. Opening the iron box: transcriptional metalloregulation by the fur protein. J Bacteriol. 1999;181:6223–6229.

26. Dumas Z, Ross-Gillespie A, Kümmerli R. Switching between apparently redundant iron-uptake mechanisms benefits bacteria in changeable environments Proc R Soc B. 2013;280:20131055.

27. Lamont IL, Beare P, Ochsner U, Vasil AI, Vasil ML. Siderophore-mediated signaling regulates virulence factor production in *Pseudomonas aeruginosa*. Proc Natl Acad Sci USA. 2002;99:7072–7077.

28. Michel L, Bachelard A, Reimmann C. Ferripyochelin uptake genes are involved in pyochelin-mediated signalling in *Pseudomonas aeruginosa*. Microbiology. 2007;153:1508–1518.

29. Cornelis P, Dingemans J. *Pseudomonas aeruginosa* adapts its iron uptake strategies in function of the type of infections. Front Cell Infect Microbiol. 2013;3:75.

30. Tiburzi F, Imperi F, Visca P. Intracellular levels and activity of PvdS, the major iron starvation sigma factor of *Pseudomonas aeruginosa*. Mol Microbiol. 2008;67:213–227.

31. Kümmerli R, Jiricny N, Clarke LS, West SA, Griffin AS. Phenotypic plasticity of a cooperative behaviour in bacteria. J Evol Biol. 2009;22:589–598.

32. Harrison F. Dynamic social behaviour in a bacterium: *Pseudomonas aeruginosa* partially compensates for siderophore loss to cheats. J Evol Biol. 2013;26:1370–1378.

33. Schiessl KT, Ross-Gillespie A, Cornforth DM, Weigert M, Bigosch C, Brown SP et al. Individual-versus group-optimality in the production of secreted bacterial compounds. Evolution. 2019;73:675–688.

34. Ross-Gillespie A, Dumas Z, Kümmerli R. Evolutionary dynamics of interlinked public goods traits: an experimental study of siderophore production in *Pseudomonas aeruginosa*. J Evol Biol. 2015;28:29–39.

35. Choi K-H, Schweizer HP. mini-Tn7 insertion in bacteria with single attTn7 sites: example *Pseudomonas aeruginosa*. Nat Protoc. 2006;1:153–161.

36. Rezzoagli C, Granato ET, Kümmerli R. In-vivo microscopy reveals the impact of *Pseudomonas aeruginosa* social interactions on host colonization. ISME J. 2019;13:2403–2414.

37. Minoia M, Gaillard M, Reinhard F, Stojanov M, Sentchilo V, van der Meer JR. Stochasticity and bistability in horizontal transfer control of a genomic island in *Pseudomonas*. Proc Natl Acad Sci USA. 2008;105:20792–20797.

38. Meyer J-M, Neely A, Stintzi A, Georges C, Holder IA. Pyoverdin is essential for viruence of *Pseudomonas aeruginosa*. Infect Immun. 1996;64:518–523.

39. de Jong IG, Beilharz K, Kuipers OP, Veening JW. Live cell imaging of Bacillus subtilis and Streptococcus pneumoniae using automated time-lapse microscopy. J Vis Exp. 2011;53:e3145.

40. Weigert M, Kümmerli R. The physical boundaries of public goods cooperation between surface-attached bacterial cells. Proc R Soc B. 2017;284:20170631.

41. Berg S, Kutra D, Kroeger T, Straehle CN, Kausler BX, Haubold C et al. ilastik: interactive machine learning for (bio)image analysis. Nat Methods. 2019;16:1226–1232.

42. Schindelin J, Arganda-Carreras I, Frise E, Kaynig V, Longair M, Pietzsch T et al. Fiji: an open-source platform for biological-image analysis. Nat Methods. 2012;9:676–682.

43. Cunrath O, Gasser V, Hoegy F, Reimmann C, Guillon L, Schalk IJ. A cell biological view of the siderophore pyochelin iron uptake pathway in *Pseudomonas aeruginosa*. Environ Microbiol. 2014;17:171–185.

44. van der Veen DR, Riede SJ, Heideman PD, Hau M, van der Vinne V, Hut RA. Flexible clock systems: adjusting the temporal programme. Phil Trans R Soc B. 2017;372.

45. Helm B, Visser ME, Schwartz W, Kronfeld-Schor N, Gerkema M, Piersma T et al. Two sides of a coin: ecological and chronobiological perspectives of timing in the wild. Phil Trans R Soc B. 2017;372.

46. Andrews SC, Robinson AK, Rodriguez-Quinones F. Bacterial iron homeostasis. FEMS Microbiol Rev. 2003;27:215–237.

47. Alqarni B, Colley B, Klebensberger J, McDougald D, Rice SA. Expression stability of 13 housekeeping genes during carbon starvation of *Pseudomonas aeruginosa*. Journal of Microbiological Methods. 2016;127:182–187.

48. Veening J-W, Smits WK, Kuipers OP. Bistability, epigenetics, and bet-hedging in bacteria. Annu Rev Microbiol. 2008;62:193–210.

49. Ratcliff WC, Denison RF. Individual-level bet hedging in the bacterium *Sinorhizobium meliloti*. Curr Biol. 2010;20:1740–1744.

50. Schreiber F, Littman S, Lavik G, Esrig S, Meiborn A, Kuypers MMM et al. Phenotypic heterogeneity driven by nutrient limitation promotes growth in fluctuating environments. Nat Microbiol. 2016;1:16055.

51. Leinweber A, Weigert M, Kümmerli R. The bacterium *Pseudomonas aeruginosa* senses and gradually responds to interspecific competition for iron. Evolution. 2018;72:1515–1528.

52. Conradt L, Roper TJ. Consensus decision making in animals. Trends Ecol Evol. 2005;20:449–456.

53. Sumpter DJT. The principles of collective animal behaviour. Phil Trans R Soc B. 2006;361:5–22.

54. Couzin ID. Collective cognition in animal groups. Trends Cogn Sci. 2009;13:36–43.

55. Bose T, Reina A, Marshall JAR. Collective decision-making. Current Opinion in Behavioral Sciences. 2017;16:30–34.

56. Dussutour A, Ma Q, Sumpter D. Phenotypic variability predicts decision accuracy in unicellular organisms. Proc R Soc B. 2019;286:2018–2825.

