## Supplementary Material for "From heterogeneity to homogeneity: coordination of siderophore gene expression among clonal cells of the bacterium *Pseudomonas aeruginosa*"

**These supplementary materials contain:**

- 8 supplementary figures
- 4 supplementary tables

**SUPPLEMENTARY FIGURES**

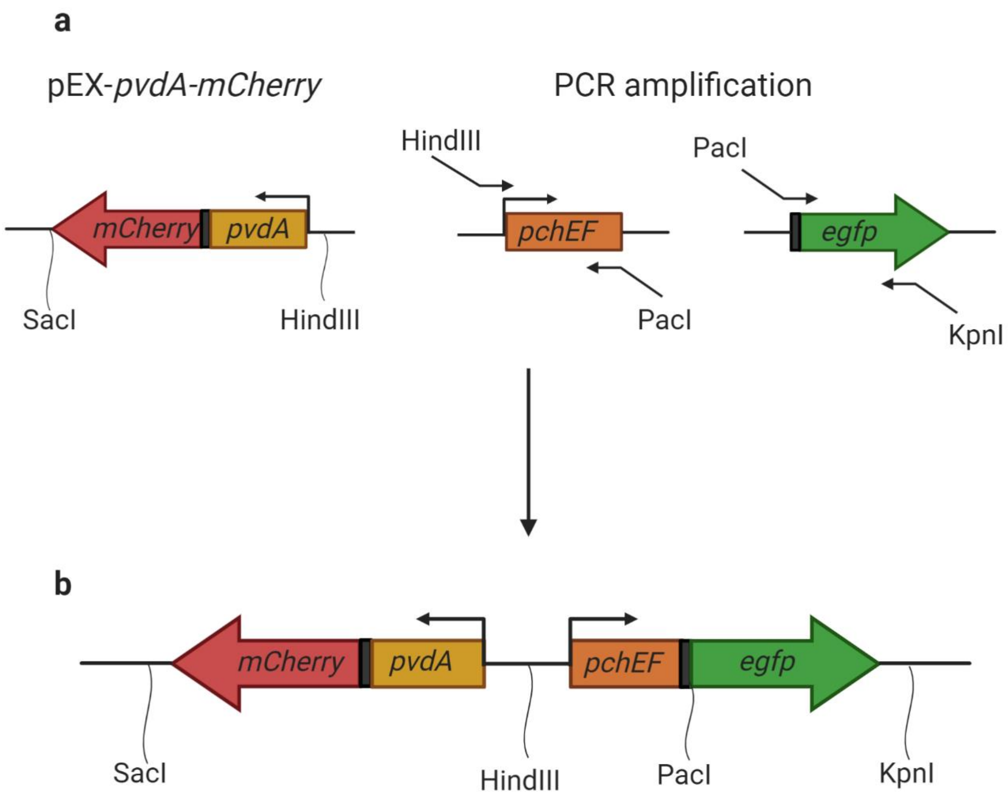

**Figure S1. Double fluorescent gene reporter scaffold type 1 used for strain PAO1*pvdA*::*mcherry–pchEF*::*egfp*. (a)** The promoters for the pyochelin synthesis genes *pchEF* and the fluorescent gene marker e*gfp* were PCR amplified using primers denoted by arrows with unique restriction enzyme sites. The vector pEX-*pvdA*-*mCherry* containing the promotor for the pyoverdine synthesis gene *pvdA* fused to the *mCherry* gene was digested at the denoted unique restriction enzyme sites. **(b)** Subsequently, the gene fragments were ligated at the specific restriction enzyme sites and integrated into the pUC18-miniTn7-Gm vector. Ribosomal binding sites are shown as dark brown rectangles at the start of the fluorescent gene markers *egfp* and *mCherry*.

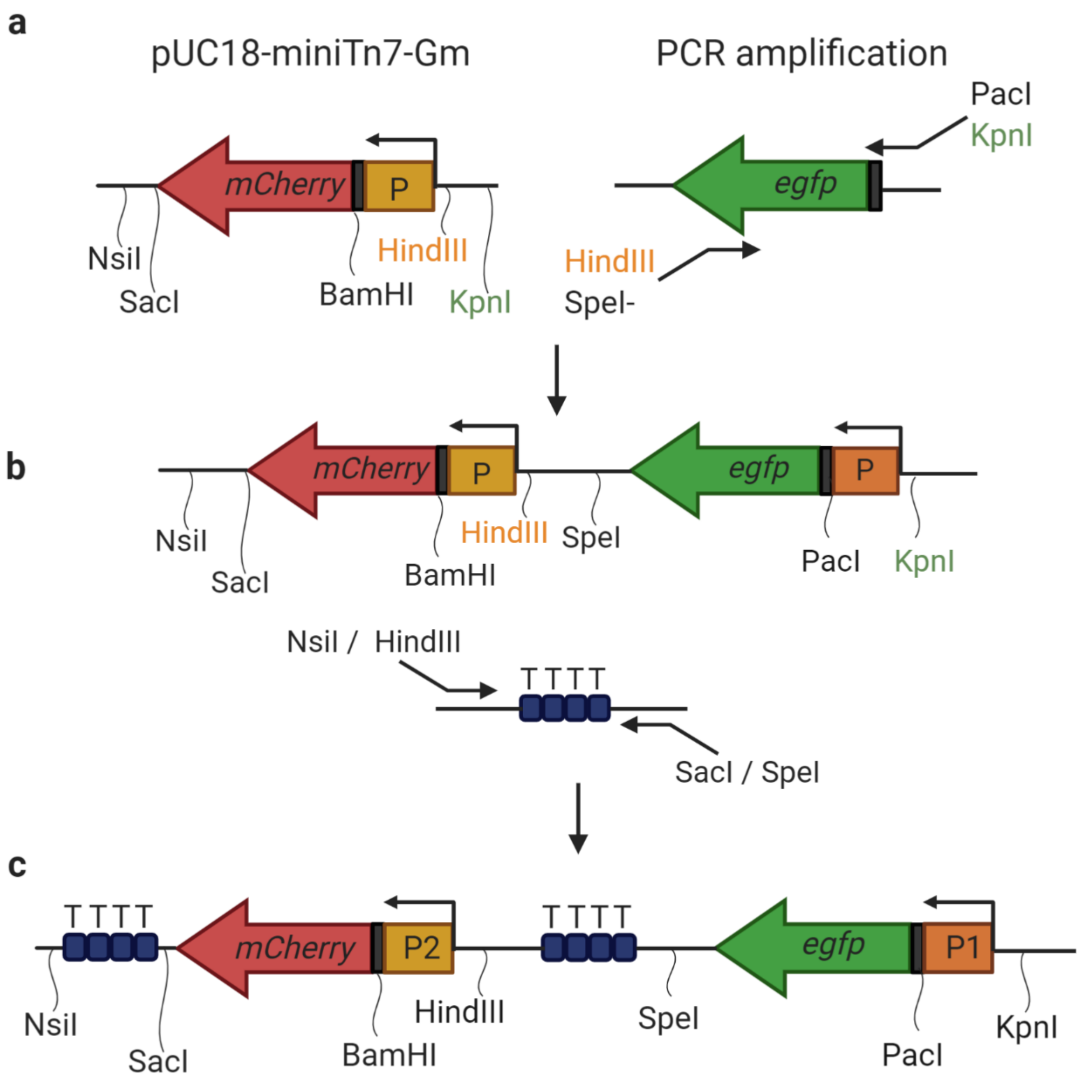

**Figure S2. Double fluorescent gene reporter scaffold type 2 used for the strains PAO1*pchEF*::*mcherry–rpsL*::*egfp* and PAO1*pvdA*::*mcherry–rpsL*::*egfp*.** **(a)**The fluorescent gene marker *egfp* was PCR amplified using the primers denoted by arrows with unique restriction enzyme sites. The PCR product contained an empty promoter region between the restriction enzyme sites PacI and KpnI. **(b)** Subsequently, the gene fragment was ligated at the specific restriction sites HindIII and KpnI (denoted by colour code of respective restriction enzyme) of the pUC18 miniTn7 Gm – *mCherry* vector plasmid containing an empty promoter site fused to *mCherry* . Four rho-independent terminators denoted by T (deep blue boxes) were also PCR amplified using two pairs of primers containing unique restrictions sites NsiI and SacI, and HindIII and SpeI. **(c)** Promoter sites are denoted as P1 (fused to *egfp*) and P2 (fused to *mCherry*). Promoter regions of the genes of interest were added at the sites P1 (fused to *egfp*)

and P2 (fused to *mCherry*) using restriction enzyme sites KpnI and PacI or HindIII and BamHI, respectively. Promoter interference was eliminated through the ligation of terminator sites between two promoter fusions at the specific restriction sites HindIII and SpeI, and NsiI and SacI. Ribosomal binding sites are shown as dark brown rectangles at the start of the fluorescent gene markers *egfp* and *mCherry*.

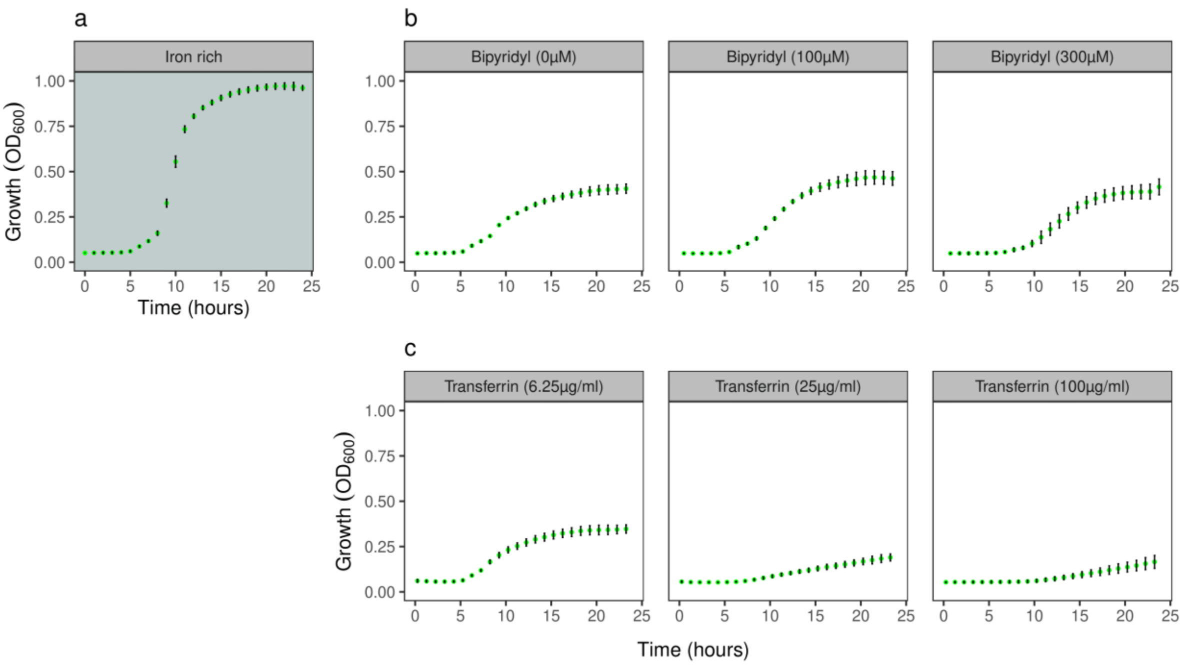

**Figure S3. Population level growth kinetics.** The panels show temporal dynamics of bacterial growth across a range of CAA media differing in their level of iron limitations. Values and error bars represent the mean growth and standard deviation across 24 replicates, in **(a)** iron-replete CAA medium (100 µM $FeCl_3$); **(b)** CAA media with increasing concentrations of the iron chelator bipyridyl; **(c)** CAA media with increasing concentrations of the iron chelator apo-transferrin. The bacterial growth is maximized in iron-replete CAA medium, and decreases with the addition of iron chelators. Note that we combined the data from three PAO1 strains (PAO1 wildtype and the two single gene reporter strains PAO1*pchEF*:*mcherry* and PAO1*pvdA*:*mcherry*) as there was no significant differences in growth between them.

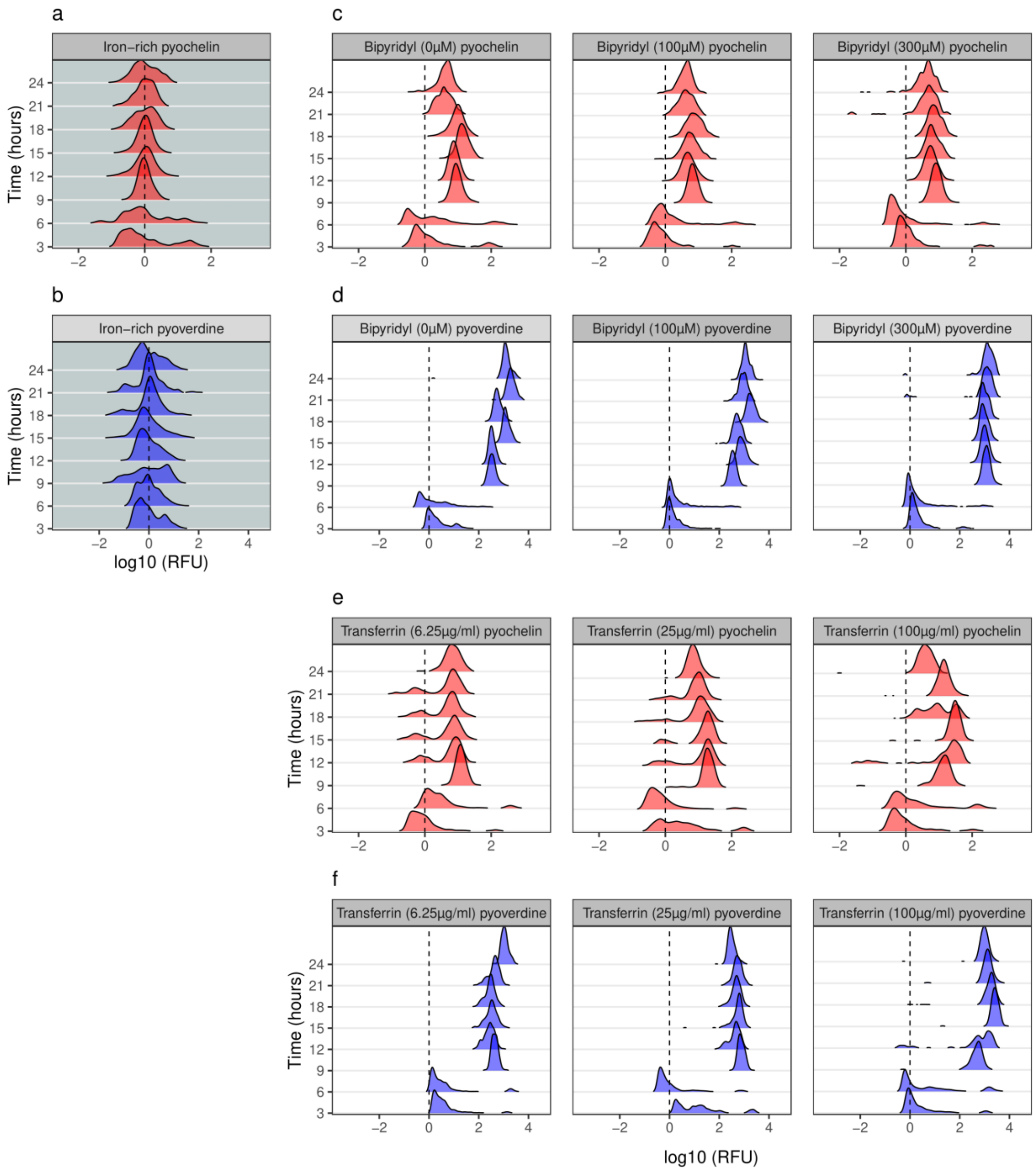

**Figure S4. Single-cell siderophore gene expression shown as density plots across time and media.** Single-cell siderophore gene expression shown as density plots for pyochelin (red: *pchEF*) and pyoverdine (blue: *pvdA)*, measured with the double reporter strain PAO1*pvdA*::*mcherry–pchEF*::*egfp* across a range of CAA media differing in their levels of iron limitations. The density plots represent the frequency distribution of log-transformed fluorescence values. **(a+b)** Iron-replete CAA medium (100 µM $FeCl_3$); **(c+d)** CAA media with increasing concentration of the iron chelator bipyridyl; **(e+f)** CAA media with increasing concentration of the iron chelator apo-transferrin. The black dashed line at 0 represents the background level of gene expression.

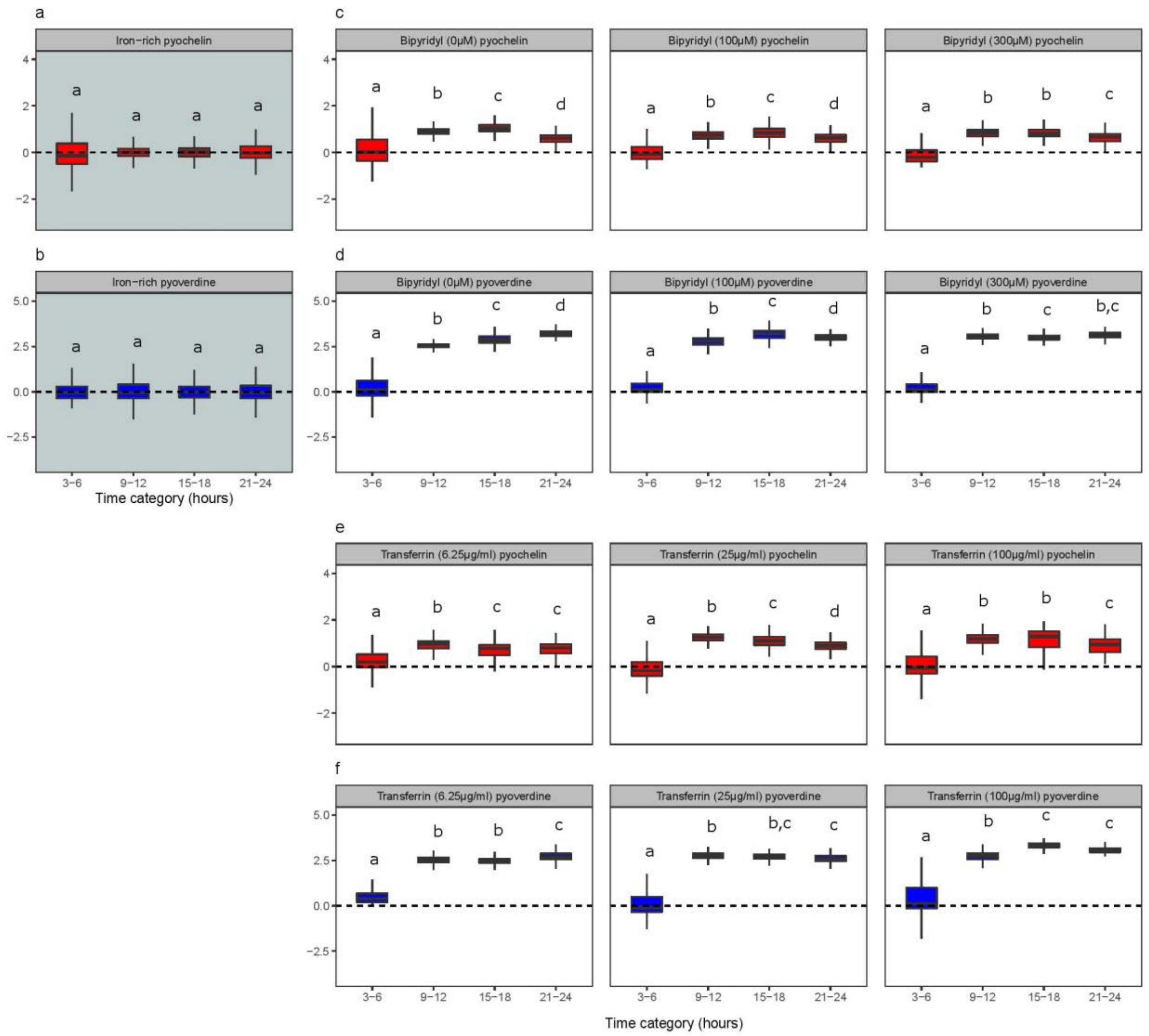

**Figure S5. Single-cell gene expression shown as box plots across time and media.** Single-cell siderophore gene expression shown as box plots for pyochelin (red: *pchEF*) and pyoverdine (blue: *pvdA)*, measured with the double gene reporter strain PAO1*pvdA*::*mcherry*–*pchEF*::*egfp* across a range of CAA media differing in their levels of iron limitations. **(a+b)** Iron-replete CAA medium (100 µM $FeCl_3$); **(c+d)** CAA media with increasing concentration of the iron chelator bipyridyl; **(e+f)** CAA media with increasing concentration of the iron chelator apo-transferrin. The boxplots show the median with the 25$^{th}$ and 75$^{th}$ percentiles of log-transformed fluorescence values for four time periods (3-6 hours; 9-12 hours; 15-18 hours; 21-24 hours). We collapsed adjacent time points because they showed similar gene expression patterns (Fig. 3). Whiskers show the 1.5 interquartile range. The black dashed line at 0 represents the background level of gene expression. The different letters above the boxplots indicate statistically significant differences between time categories.

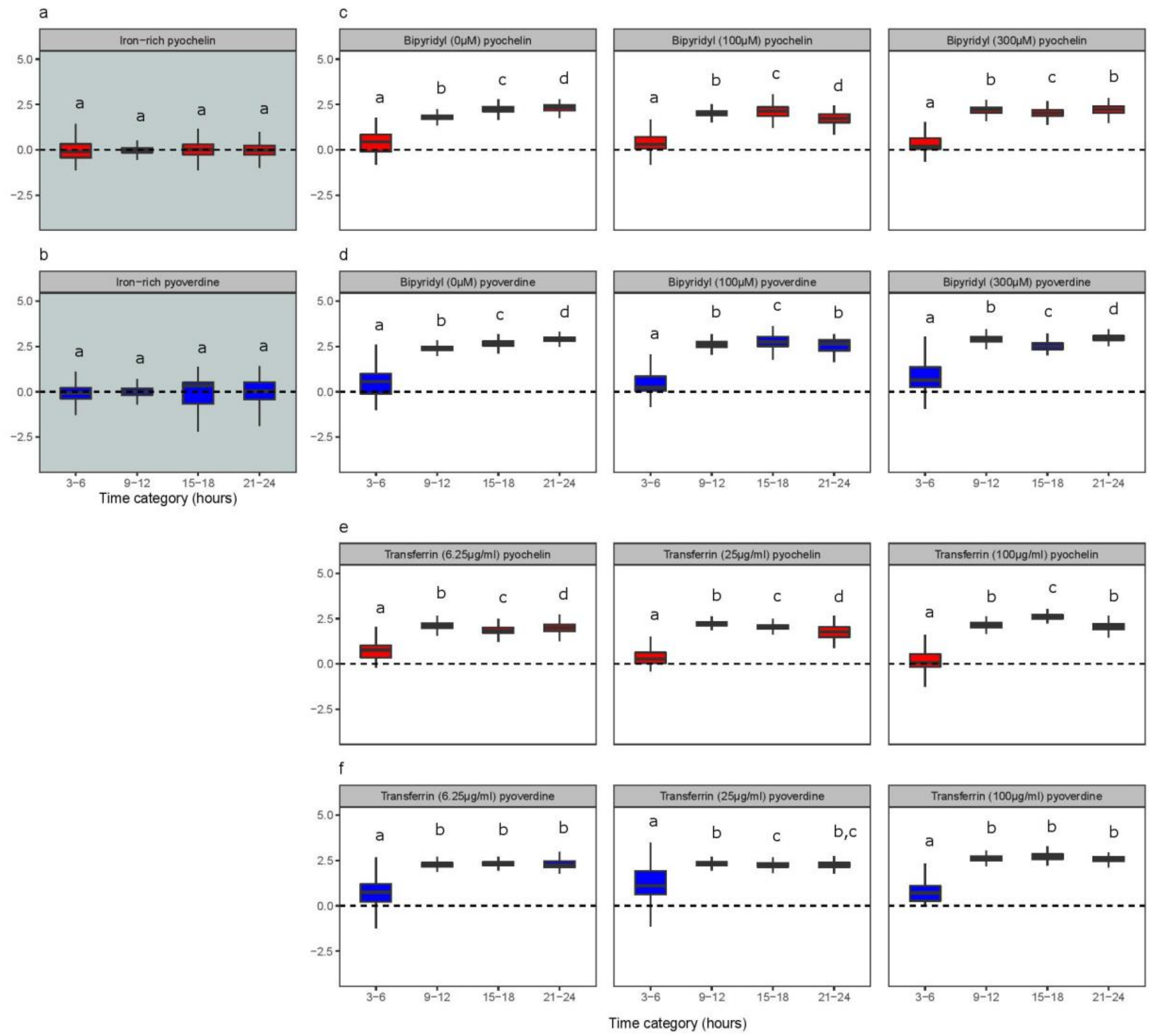

**Figure S6. Single-cell gene expression shown as box plots across time and media.** Single-cell siderophore gene expression shown as box plots for pyochelin (red: *pchEF*) and pyoverdine (blue: *pvdA)*, measured with the single gene reporters PAO1*pchEF*:*mcherry* and PAO1*pvdA*:*mcherry*, respectively across a range of CAA media differing in their levels of iron limitations. **(a+b)** Iron-replete CAA medium (100 µM $FeCl_3$); **(c+d)** CAA media with increasing concentration of the iron chelator bipyridyl; **(e+f)** CAA media with increasing concentration of the iron chelator apo-transferrin. The boxplots show the median with the 25$^{th}$ and 75$^{th}$ percentiles of log-transformed fluorescence values for four time periods (3-6 hours; 9-12 hours; 15-18 hours; 21-24 hours). We collapsed adjacent time points because they showed similar gene expression patterns. Whiskers show the 1.5 interquartile range. The black dashed line at 0 represents the background level of gene expression. The different letters above the boxplots indicate statistically significant differences between time categories.

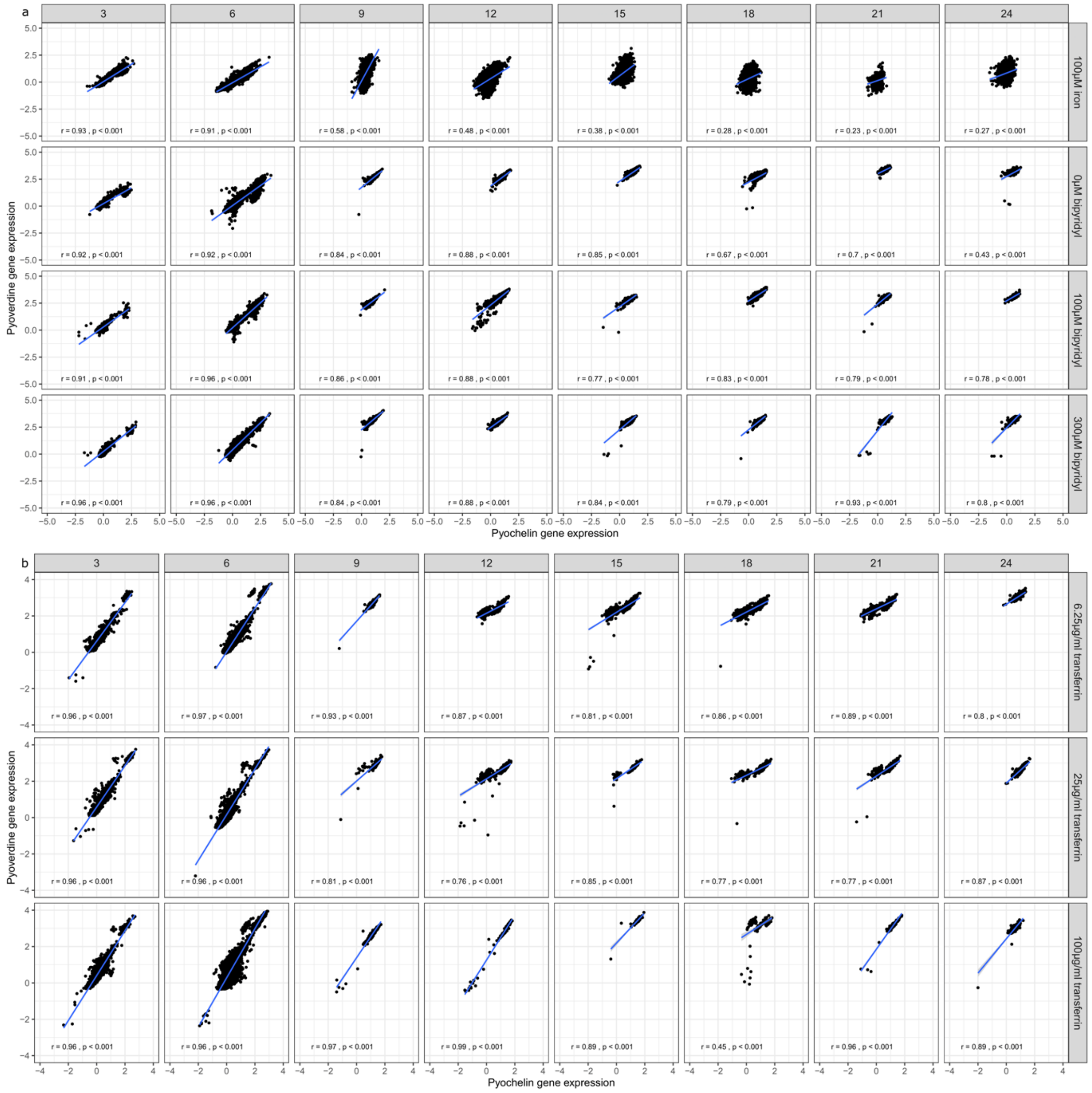

**Figure S7. Correlations between pyochelin and pyoverdine gene expression across individual cells over time and in media differing in their level of iron limitation.** All correlations between pyochelin and pyoverdine gene expression are measured with the double reporter strain PAO1*pvdA*::*mcherry–pchEF*::*egfp*. Each dot represents a single cell with its corresponding pyochelin (*pchEF)* and pyoverdine (*pvdA)* gene expression. Gene expression was measured every three hours (from left to right) by analysing a subset of cells extracted from growing cultures. Rows depict the different media conditions. **(a)** Gene expression correlations in iron-supplemented CAA medium (top row) and in CAA media supplemented with increasing concentration of the iron chelator bipyridyl (top-down). **(b)** Gene expression correlations in CAA media supplemented with increasing concentrations of the iron chelator apo-transferrin (top-down). The Pearson correlation coefficient r and the p-value are provided in each panel.

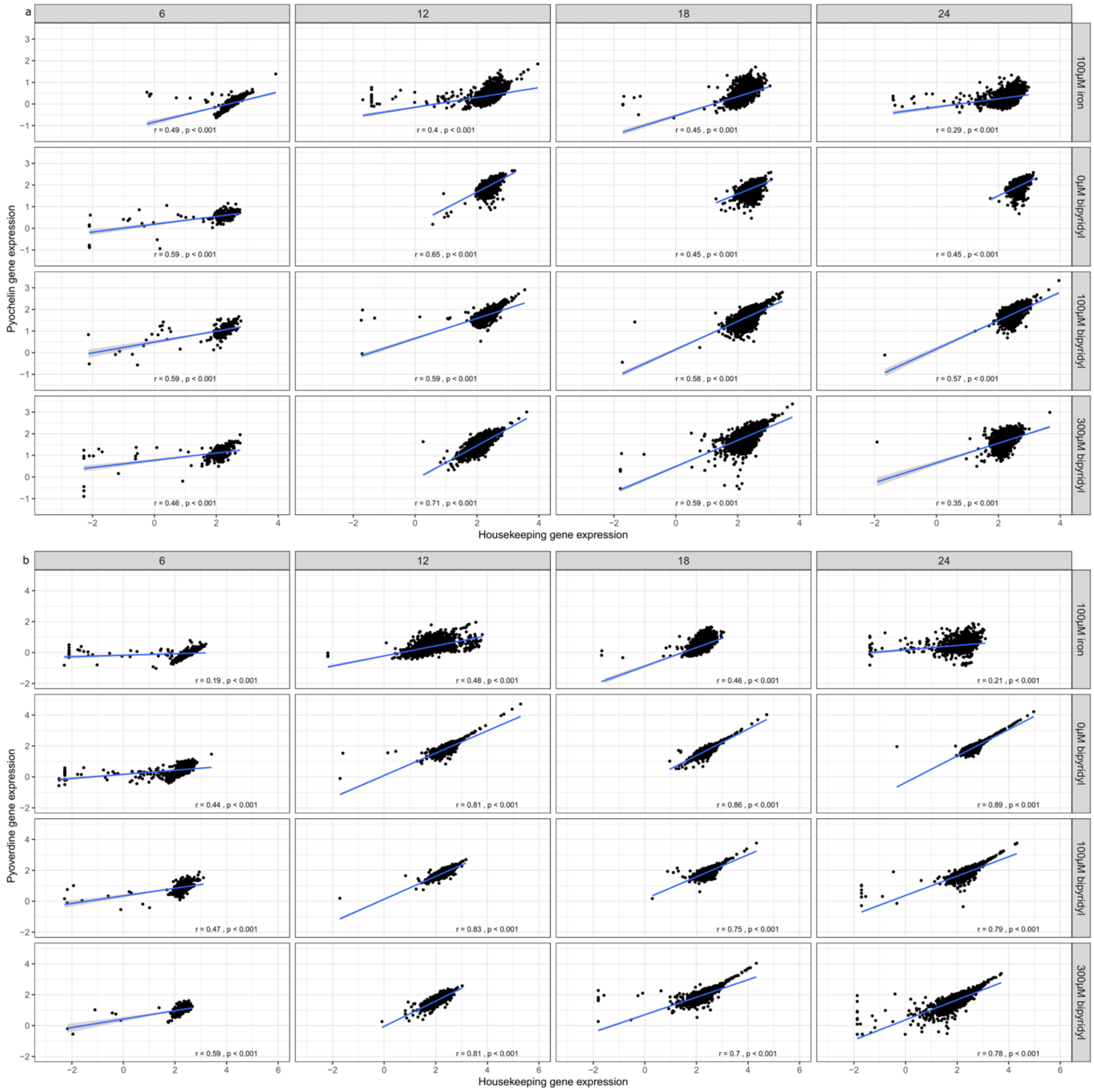

**Figure S8. Correlations between siderophore and *rpsL* housekeeping gene expression across individual cells over time and in media differing in their level of iron limitation. (a)** Correlations between pyochelin and housekeeping gene expression are measured with the double reporter strain PAO1*pchEF*::*mcherry–rpsL*::*egfp* . **(b)** Correlations between pyoverdine and housekeeping gene expression are measured with the double reporter strain PAO1*pvdA*::*mcherry–rpsL*::*egfp*. For both strains, gene expression was measured every six hours (from left to right) by analysing a subset of cells extracted from growing cultures. Rows depict the different media conditions: iron-supplemented CAA medium (top row) and CAA media supplemented with increasing concentrations of the iron chelator bipyridyl. Each dot represents a single cell. The Pearson correlation coefficient r and the p-value are provided in each panel.

**Table S1:** Comparison of growth parameters of *P. aeruginosa* populations across a range of casamino acids medium (CAA) compositions.

| CAA supplement | concentration | growth integral | growth rate (∆OD600/h) | lag phase (h) |
|---|---|---|---|---|
| FeCl3 (µM) | 100 | 12.296 ± 0.153 | 0.273 ± 0.005 | 8.087 ± 0.030 |
| No supplement | na | 4.512 ± 0.027 | 0.089 ± 0.010 | 6.761 ± 0.030 |
| Bipyridyl (µM) | 50 | 4.946 ± 0.084 | 0.132 ± 0.067 | 6.784 ± 0.134 |
| Bipyridyl (µM) | 100 | 4.993 ± 0.079 | 0.064 ± 0.001 | 7.200 ± 0.098 |
| Bipyridyl (µM) | 150 | 4.868 ± 0.066 | 0.101 ± 0.035 | 7.425 ± 0.112 |
| Bipyridyl (µM) | 200 | 4.623 ± 0.090 | 0.073 ± 0.006 | 7.492 ± 0.354 |
| Bipyridyl (µM) | 300 | 3.557 ± 0.116 | 0.051 ± 0.001 | 9.016 ± 0.154 |
| Transferrin (µg/ml) | 6.25 | 4.095 ± 0.074 | 0.082 ± 0.023 | 5.077 ± 0.401 |
| Transferrin (µg/ml) | 12.5 | 1.319 ± 0.043 | 0.015 ± 0.001 | 11.029 ± 0.696 |
| Transferrin (µg/ml) | 25 | 0.960 ± 0.021 | 0.017 ± 0.003 | 12.453 ± 0.716 |
| Transferrin (µg/ml) | 50 | 0.876 ± 0.055 | 0.016 ± 0.001 | 13.743 ± 0.756 |
| Transferrin (µg/ml) | 100 | 0.952 ± 0.028 | 0.015 ± 0.001 | 12.940 ± 0.605 |

Table S2: List of strains

| **Strain name** | **Description or genotype** | **Source or reference** |
|---|---|---|
| ***E. coli*** | | |
| CC118 λpir | Δ (*ara, leu*)$_{7697}$ *araD*139 *ΔlacX74 galE galK phoA20 thi-1 rpsE rpoB* ($Rf^R$) *argE(am) recA1 λpir*$^+$ | De Lorenzo *et al.*, 1990(1) |
| ***P. aeruginosa*** | | |
| PAO1 (ATCC 15692) | Wild type strain | This laboratory |
| ***P. aeruginosa* PAO1 (single fluorescent gene reporters)** | | |
| PAO1*pvdA:mcherry* | Transcriptional fusion *pvdA::mcherry* from pSR01 | Rezzoagli *et al.* 2019(2) |
| PAO1*pchEF:mcherry* | Transcriptional fusion *pchEF::mcherry* from pSR02 | Rezzoagli *et al.* 2019(2) |
| PAO1*rpsL:*mcherry | Transcriptional fusion *rpsL*::mcherry from pSR03 | Jayakumar *et al.* (unpublished, this laboratory) |
| ***P. aeruginosa* PAO1 (double fluorescent gene reporters)** | | |
| PAO1*pvdA*: mcherry-*pchEF*::GFP | Transcriptional fusion *pvdA*::mcherry and *pchEF*::GFP from pDR01 | This study |
| PAO1*pvdA*: mcherry-*rpsL*::GFP | Transcriptional fusion *pvdA*::mcherry and *rpsL*::GFP from pDR02 | This study |
| PAO1*pchEF*: mcherry-*rpsL*::GFP | Transcriptional fusion *pchEF*::mcherry and *rpsL*::GFP from pDR03 | This study |

Table S3: List of plasmids

| Plasmid name | Description or genotype | Source or reference |
|---|---|---|
| pEX-A128-*pvdA::mcherry* | Commercial plasmid with *pchEF::mcherry* between HindIII/SacI sites | Weigert *et al*., 2017(3) |
| pUX-BF13 | Helper plasmid to provide Tn7 transposase proteins | Bao *et al*., 1991(4) |
| pUC18-miniTn7-Gm | $Gm^r$ on mini-Tn7; for chromosomal insertion in $Gm^s$ bacteria in the *att*Tn7 site | Choi and Schweizer *et al*., 2006(5) |
| pUC18-miniTn7-Gm-mcherry-GFP | Derived from pUC18-miniTn7-Gm-mcherry; with amplified GFP from pEX-A128-*pchEF*::GFP | This study |
| pDR01 | pUC18-mini-Tn7-Gm with *pvdA*::mcherry and *pchEF*::GFP | This study |
| pDR02 | pUC18-mini-Tn7-Gm with *pvdA*::mcherry and *rpsL*::GFP | This study |
| pDR03 | pUC18-mini-Tn7-Gm with *pchEF*::mcherry and *rpsL*::GFP | This study |

Table S4: List of primers

| **Primer name** | **Sequence (5'-3')** | **Application** | **Template** |
|---|---|---|---|
| pvdA_Rev_Sac1 | CGG CAT CAG AGC AGA TTG TA | Plasmid pDR01 | pEX-pvdA-*mcherry* |
| pvdA_Fwd_HindIII | GGA TCC AAG CGA GCA AAA G | Plasmid pDR01 | pEX-pvdA-*mcherry* |
| pchEF_Fwd_HindIII | GATCAA GCTTCAAGCGCTACG GCATCTC | Plasmid pDR01 & pDR03 | PAO1 gDNA |
| pchEF_Rev_PacI | CGA G TTAATTAA TC ACT GCT CGG TCA GCC AGT C | Plasmid pDR01 | PAO1 gDNA |
| pvdA_Fwd_HindIII_2 | CAGTGCAGGTGGGAAGCTTATGC | Plasmid pDR02 | pEX-pvdA-*mcherry* |
| pvdA_Rev_BamHI | CAGTCCTCCTTCTTAAAGGGATCC | Plasmid pDR02 | pEX-pvdA-*mcherry* |
| rpsL_Fwd_HindIII | CAGTAAGCTTGTACCGGTCTGGCTTAC CAC | Plasmid pDR02 & pDR03 | PAO1 gDNA |
| rpsL_Rev_BamHI | CAGTGGATCCTCAGTGTGCCGAGTTCG GCTTTT | Plasmid pDR02 & pDR03 | PAO1 gDNA |
| pchEF_Rev_BamHI | CGA GGGATCCTC ACTGCTCGGTCAGCCAGT C | Plasmid pDR03 | PAO1 gDNA |
| Fwd_HindIII_Tn7 (for pDR02) | GCGCGAATGGGAAGCCGACTG | *E. coli* colonies | Colony PCR to check insertion in *E. coli* |
| Fwd_HindIII_pEx (pDR03) | GAT CAA GCT TCA AGC GCT ACG GCA TCT C | *E. coli* colonies | Colony PCR to |

| | | | |
|---|---|---|---|
| | | | check insertion in *E. coli* |
| Rev_Tn7_primer (for pDR02 & pDR03) | CGAACCGAACAGGCTTATGT | *E. coli* colonies | Colony PCR to check insertion in *E. coli* |
| mcherry_rev2 | GGATATCCGCTGGGTGTTTA | *P. aeruginosa* colonies | Colony PCR to check insertion in *P. aeruginosa* |

To confirm the insertion of the promoter region into the miniTn7 vector, colony PCR was performed using the mcherry_rev2 primer, together with the corresponding "Fwd_HindIII" promoter-specific primers

References

1. V. De Lorenzo, M. Herrero, U. Jakubzik, K. N. Timmis, Mini-Tn5 transposon derivatives for insertion mutagenesis, promoter probing, and chromosomal insertion of cloned DNA in gram-negative eubacteria. *J. Bacteriol.* **172**, 6568–6572 (1990).

2. C. Rezzoagli, E. T. Granato, R. Kümmerli, In-vivo microscopy reveals the impact of Pseudomonas aeruginosa social interactions on host colonization. *ISME J.* (2019) https:/doi.org/10.1038/s41396-019-0442-8.

3. M. Weigert, R. Kümmerli, The physical boundaries of public goods cooperation between surface-attached bacterial cells. *Proc. R. Soc. B Biol. Sci.* **284** (2017).

4. Y. Bao, D. P. Lies, H. Fu, G. P. Roberts, An improved Tn7-based system for the single-copy insertion of cloned genes into chromosomes of gram-negative bacteria. *Gene* **109**, 167–168 (1991).

5. K. H. Choi, H. P. Schweizer, mini-Tn7 insertion in bacteria with single attTn7 sites: Example Pseudomonas aeruginosa. *Nat. Protoc.* **1**, 153–161 (2006).
